# Automating Candidate Gene Prioritization with Large Language Models: From Naive Scoring to Literature-Grounded Validation

**DOI:** 10.1101/2025.09.17.676837

**Authors:** Taushif Khan, Mohammed Toufiq, Marina Yurieva, Nitaya Indrawattana, Akanitt Jittmittraphap, Nathamon Kosoltanapiwat, Pornpan Pumirat, Passanesh Sukphopetch, Muthita Vanaporn, Karolina Palucka, Basirudeen Kabeer, Darawan Rinchai, Damien Chaussabel

## Abstract

**Background:** Identifying promising therapeutic targets from thousands of genes in transcriptomic studies remains a major bottleneck in biomedical research. While large language models (LLMs) show potential for gene prioritization, they suffer from hallucination and lack systematic validation against expert knowledge.

**Methods:** We developed a two-stage computational framework that combines LLM-based screening with literature validation for systematic gene prioritization. Starting with 10,824 genes from the BloodGen3 repertoire, we applied multi-criteria evaluation for sepsis relevance, followed by retrieval-augmented generation (RAG) using 6,346 curated sepsis publications. A novel faithfulness evaluation system verified that LLM predictions aligned with retrieved literature evidence.

**Results:** The framework identified 609 sepsis-relevant genes with >94% filtering efficiency, demonstrating strong enrichment for inflammatory pathways including TNF-α signaling, complement activation, and interferon responses. Literature validation yielded 30 ultra-high confidence therapeutic candidates, including both established sepsis genes (IL10, TREM1, S100A9, NLRP3) and novel targets warranting investigation. Benchmark validation against expert-curated databases achieved 71.2% recall, with systematic correlation between computational confidence and evidence quality. The final candidate set balanced discovery (11 novel genes) with validation (19 known genes), maintaining biological coherence throughout the filtering process.

**Conclusions:** This framework demonstrates that rigorous methodology can transform unreliable LLM outputs into systematically validated biological insights. By combining computational efficiency with literature grounding, the approach provides a practical tool for prioritizing experimental validation efforts. The modular design enables adaptation to other diseases through knowledge base substitution, offering a systematic approach to literature-guided biomarker discovery.

**Availability:** Source code and implementation details are available at https://github.com/taushifkhan/llm-geneprioritization-framework, vector database at https://doi.org/10.5281/zenodo.15802241 and Interactive demonstration at https://llm-geneprioritization.streamlit.app/

## 1. INTRODUCTION

Candidate gene prioritization plays a crucial role in identifying potential biomarkers from large-scale molecular profiling data. Systems-scale profiling technologies, such as transcriptomics, have revolutionized biomedical research by simultaneously measuring tens of thousands of analytes, leading to significant advances in oncology (1,2), autoimmunity (3,4), and infectious diseases (5,6). However, translating these findings into actionable clinical insights requires identifying relevant analyte panels and designing targeted profiling assays (7,8). Targeted transcriptional profiling assays enable precise, quantitative assessments of tens to hundreds of transcripts (9,10), offering cost-effectiveness, rapid turnaround times, and high-throughput capability (11–13). The critical challenge lies in selecting relevant candidate genes from extensive biomedical information volumes generated by systems-scale profiling technologies.

Traditional knowledge-driven methods face multiple challenges in efficiently processing vast literature to identify promising candidates. While gene ontologies and curated pathways provide valuable information, they rely on static knowledge bases that may not capture current research findings or miss relevant associations buried within literature (14). Moreover, traditional approaches rely heavily on static knowledge bases that may not capture the most current research findings or may miss relevant associations buried within the literature (15). Manual curation, though thorough, is time-intensive and may lack comprehensive coverage due to information volume constraints.

Computational approaches have emerged to address these limitations. Methods like ProGENI integrate protein-protein and genetic interactions with expression data for drug response prediction(16), while the Monarch Initiative combines multi-species data for variant prioritization(17). However, recent large language model (LLM) applications in biomedical gene prioritization show mixed results. Kim et al. achieved only 16.0% accuracy with GPT-4 for phenotype-driven gene prioritization in rare genetic disorders, lagging traditional bioinformatics tools (18). These limitations highlight the need for sophisticated approaches addressing fundamental challenges in LLM-based biomedical inference.

LLMs such as GPT-4, Claude, and PaLM have demonstrated remarkable natural language understanding capabilities (15,19,20), trained on vast text data enabling synthesis of diverse information sources including scientific literature. However, naive LLM approaches face significant biomedical limitations: potential hallucinations, reliance on training data snapshots potentially missing current knowledge, and difficulty providing verifiable, literature-grounded justifications.

Recent advances in retrieval-augmented generation (RAG) and chain-of-thought reasoning offer promising solutions (21). RAG systems dynamically retrieve and incorporate relevant literature, providing current and verifiable evidence for gene prioritization. Combined with faithfulness evaluation techniques verifying model outputs against retrieved evidence, these approaches significantly enhance reliability and interpretability. Chain-of-Thought reasoning enables multi-perspective evaluation and structured synthesis of competing information sources.

We previously demonstrated LLM utility in manual candidate gene prioritization (22,23), focusing on circulating erythroid cell blood transcriptional signatures associated with respiratory syncytial virus disease severity (5), vaccine response (6), and metastatic melanoma(5). Benchmarking four LLMs across multiple criteria, GPT-4 and Claude showed superior performance through consistent scoring (correlation coefficients > 0.8), strong alignment with manual literature curation, and evidence-based justifications. This established foundations for automated gene prioritization while highlighting needs for systematic validation and evidence verification (22,24).

Building upon this work, we developed a comprehensive framework integrating multiple advanced NLP techniques to address naive LLM limitations. Our framework combines RAG using curated biomedical literature, systematic faithfulness evaluation reducing hallucinations, and Chain-of-Thought reasoning for structured evidence synthesis. This multi-layered approach enables literature-grounded gene prioritization with systematic validation against external reference standards and functional biological coherence assessment, addressing key challenges: verifiable evidence grounding, systematic uncertainty handling, and scalable processing while maintaining interpretability.

This integration of advanced NLP techniques into a validated candidate gene prioritization pipeline represents significant methodological advancement beyond routine LLM applications. The framework enables systematic processing of extensive module repertoires like BloodGen3 (10) while providing robust validation through external benchmarking, cross-model comparison, and biological coherence assessment, accelerating translation of systems-scale profiling data into validated, literature-supported targeted assays for clinical and research applications.

## 2. METHODS

### 2.1. Automated Gene Prioritization Framework

We developed a comprehensive workflow accessing large language models for automated gene prioritization using LlamaIndex (v0.12.37) for RAG development. Initial naive LLM runs were conducted using GPT-4T (April 2024) and GPT-4o (April 2025), with GPT-4o subsequently serving as RAG synthesizer and final arbitrator. The framework incorporates retrieval-augmented generation, faithfulness evaluation, and Chain-of-Thought reasoning, with Ollama (v0.4.8) deployment for local access to open-source models (particularly Phi-4 for faithfulness evaluation). Biomedical named entity recognition used SpaCy (v3.7.5) (25). The system handles prompt generation, model communication, response processing, and data storage in an integrated pipeline using Python 3.8 and OpenAI API (v1.75). A flow chart of the pipeline is given in **Figure 1**.

**Figure 1.**
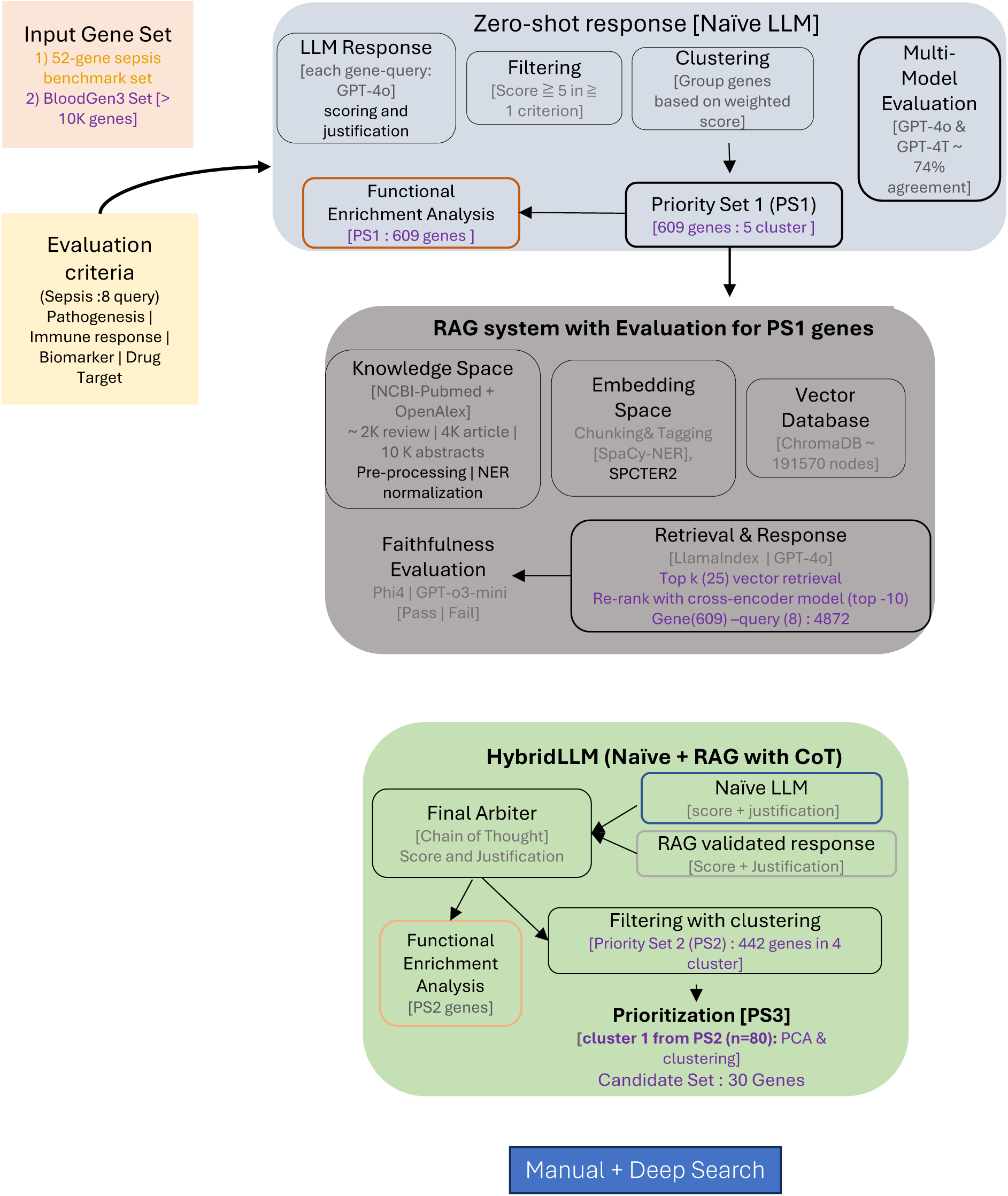
Computational Framework for Sepsis Gene Prioritization: Multi-Stage LLM-Based Evaluation with Literature Validation and Multi-Dimensional Optimization Workflow Overview: The framework processes input gene sets through progressive filtering and validation stages to identify high-confidence sepsis therapeutic targets. The workflow was first run on the 52-gene sepsis benchmark set (results in Fig 2) before being applied to the full BloodGen3 compendium. **Stage 1 (Gray):** Zero-shot evaluation using Naive LLM across eight sepsis-related criteria (pathogenesis association, immune response, biomarker potential, drug target feasibility) applied to BloodGen3 gene set (>10K genes). Genes meeting scoring thresholds undergo clustering and multi-modal evaluation, generating Priority Set 1 (PS1, 609 genes). **Stage 2 (Teal):** RAG-enhanced evaluation system processes PS1 genes through sepsis-specific knowledge base (∼6K literature sources, >10K abstracts) using SPECTER2 embeddings and LlamaIndex framework. Retrieved literature undergoes faithfulness evaluation, and RAG-validated responses combine with Naive LLM outputs in HybridLLM approach using chain-of-thought reasoning. **Stage 3 (Green):** Literature-validated genes (Priority Set 2, PS2, 442 genes) undergo functional enrichment analysis and multi-dimensional clustering. PCA-based optimization identifies top-performing gene cluster, generating ultra-high confidence Priority Set 3 (PS3, 82 genes) and candidate set (30 genes) . **Final Validation (Blue):** PS3 candidates undergo manual curation and deep-search verification to ensure robust evidence support. Color coding distinguishes processing stages: computational evaluation (gray), literature validation (teal), functional optimization (green), and manual verification (blue). The progressive filtering approach reduces 10,824 genes to 30 high-confidence therapeutic targets through systematic computational assessment, literature grounding, and multi-dimensional optimization.

### 2.2. Gene Scoring and Prompt Design

Prompts elicited gene information including official name, function summary, and quantitative scores (0-10 scale) across biomedically relevant criteria. The scoring rubric: 0 = no evidence; 1-3 = very limited evidence; 4-6 = some evidence requiring validation; 7-8 = good evidence; 9-10 = strong evidence. For sepsis prioritization, we assessed eight criteria: pathogenesis association, host immune response, organ dysfunction, circulating leukocyte biology, clinical biomarker use, blood transcriptional biomarker potential, drug target status, and therapeutic relevance. Complete templates are in Supplementary Methods (Section 3).

### 2.3. RAG Pipeline Implementation

#### Knowledge Base Construction

We curated 6,346 sepsis-related documents (4,441 articles, 1,905 reviews, 1990-2025) from NCBI PubMed Open Access, filtered using OpenAlex for high citation percentile (>0.8) (Figure S1). To enhance coverage, we supplemented this collection with 9,557 abstracts from relevant sepsis publications that were behind paywalls, ensuring comprehensive literature representation across both freely accessible and subscription-based sources. Documents underwent preprocessing (reference removal, chunk segmentation, biomedical NER), SPECTER2 embedding (allenai/specter2_base), and ChromaDB storage with metadata tagging. Detail of knowledge set curation and vector database is in Supplementary Methods (Section 1).

#### Retrieval and Synthesis

For each gene-query instance, we implemented two-stage retrieval: vector similarity search (top_k=25) followed by cross-encoder reranking (SentenceTransformerRerank, top_k=10). Retrieved context was formatted into structured prompts for GPT-4o (temperature=0.1) with explicit source attribution, maintaining query isolation to prevent cross-contamination.

#### Faithfulness Evaluation

We conducted comparative faithfulness analysis using two independent evaluators (Phi-4 and GPT-o3-mini) on a representative set of 2,928 gene-query instances from 399 genes (Figure S2). Each GPT-4o justification received binary classification (“Pass”/“Fail”) from both evaluators. Dual-evaluator agreement reached 71.94% overall with 90.6% agreement in high-confidence cases (Table S1, Figure S3). Phi-4 was selected as primary evaluator based on: (1) architectural independence from GPT-4o reducing bias, (2) consistent alignment with GPT-o3-mini in strong-evidence cases, (3) lower variance in ambiguous cases, and (4) local deployment via Ollama enabling cost-effective, reproducible evaluation. Only faithfulness-passing instances proceeded to hybrid evaluation. See Supplementary Methods on detail of evaluation (Section 2).

#### Chain-of-Thought Hybrid Reasoning

We developed a hybrid evaluation strategy synthesizing naive LLM knowledge (GPT-4o) with retrieved contextual evidence (RAG) to resolve inference divergences and persistent ambiguities. The Chain-of-Thought framework employed three structured roles: (1) Naive LLM Critic assessing initial predictions for assumptions and overconfidence, (2) Retrieved Evidence Analyst evaluating contextual support quality and specificity, and (3) Final Arbiter synthesizing perspectives with preference for strong retrieved evidence and explicit reasoning for discrepancies. This produced unified outputs: decision classification (High/Medium/Low), recalibrated score (0–10), and detailed scientific explanation. All used prompts in this work have been provided in the Supplementary Methods (Section 3).

### 2.4. Framework Validation Through Benchmark Analysis

#### Benchmark Dataset Selection and Validation Strategy

To establish framework validity before large-scale application, we systematically evaluated performance against curated sepsis gene sets from two complementary databases. DisGeNET (n=32)(26) was selected for mechanistic gene-disease associations derived from systematic literature curation with transparent evidence scoring (ScoreGDA), while CTD (n=48)(27) provided therapeutic and chemical interaction relationships with experimental validation emphasis. These databases were chosen based on methodological transparency, evidence quality standards, and peer-reviewed curation processes (Supplementary Methods: “Benchmarking Strategy and Dataset Selection”). The combined gene sets yielded 52 unique genes with established sepsis associations through expert curation (referred to as the “52-gene sepsis benchmark set”). Additionally, we have compiled a reference gene set (n=929 genes) from other known datasets (like Entrez: DisGeNET (28), CTD, Monarch (17), OpenTarget (29), Sepon and IPA(30)) (Figure S4). Detail of gene set curation for evaluation is given in Supplementary Methods (section 4).

### 2.5. Gene-Specific Weighted Scoring Calculation

Each gene received scores (0-10 scale) across eight sepsis-related evaluation criteria, as mentioned above (Figure S5). Individual criterion scores were categorized into three confidence bins: High (≥7), Medium (4–6), and Low (≤3). For each gene, the proportion of scores in each bin was calculated by dividing the count by the total number of criteria (n= 8). Weighted scores were computed using confidence-based weights: High × 1.0, Medium × 0.7, and Low × 0.3, with final scores ranging from 0.0 to 1.0. This weighted aggregation was essential to address the inherent unreliability of individual LLM scores by strategically rewarding genes that demonstrated consistent high-confidence evidence across multiple independent evaluation criteria. This approach helps in prioritizing robust multi-dimensional relevance over potentially misleading individual assessment. This methodology was applied consistently across naive LLM, RAG, and hybrid evaluation approaches to enable direct cross-method comparison.

### 2.6. Statistical Analysis and Performance Evaluation

Performance assessment used contingency tables for score transition analysis and classification reports computing precision, recall, and F1-scores. Correlation coefficients were calculated for continuous comparisons with statistical significance testing where appropriate.

## 3. RESULTS

Our aim was to convert large-language-model (LLM) outputs into a rigorously validated list of sepsis-relevant genes. To do so, we built a three-stage pipeline that (i) assigns composite scores to every gene with a naïve LLM, (ii) filters those scores through a retrieval-augmented generation (RAG) step with dual-model faithfulness checks, and (iii) refines the surviving genes by multi-dimensional optimization (Figure 1). We first asked whether this framework could recover genes that expert curators already accept as sepsis-associated before deploying it at genome scale.

### 3.1. Benchmark-Based Validation of the Framework

To establish framework validity before large-scale application, we evaluated performance against a 52-gene sepsis benchmark set derived from DisGeNET (n=32 genes) and CTD (n=48 genes) representing mechanistic and therapeutic associations respectively. The framework achieved 71.2% recall (37/52 genes) using our scoring threshold of ≥5 in any evaluation category, with systematic performance variation between databases: DisGeNET 84.4% recall (27/32) and CTD 70.8% recall (34/48) (Table S2). Analysis revealed systematic correlation with expert evidence quality, with identified genes showing significantly higher curation scores than missed genes (Figure 2A). The three evaluation approaches exhibited distinct but appropriate scoring behaviors (Figure 2B). All methods demonstrated expected correlations with PubMed publication frequency that mirror expert-curated databases: naive LLM showed strong correlation (Spearman ρ = 0.795, p < 0.001), while RAG and hybrid approaches showed moderate correlations (ρ = 0.528 and ρ = 0.524 respectively, both p < 0.001) (Table S3). These correlations align with benchmark database patterns (ScoreGDA ρ = 0.802, CTD ρ = 0.515), confirming that higher scores appropriately reflect genes with more robust evidence bases rather than random scoring behavior. Notably, RAG showed specific enhancements for less-studied genes (AQP1, MIR483), suggesting capability to identify promising but understudied candidates. (Table S2).

**Figure 2.**
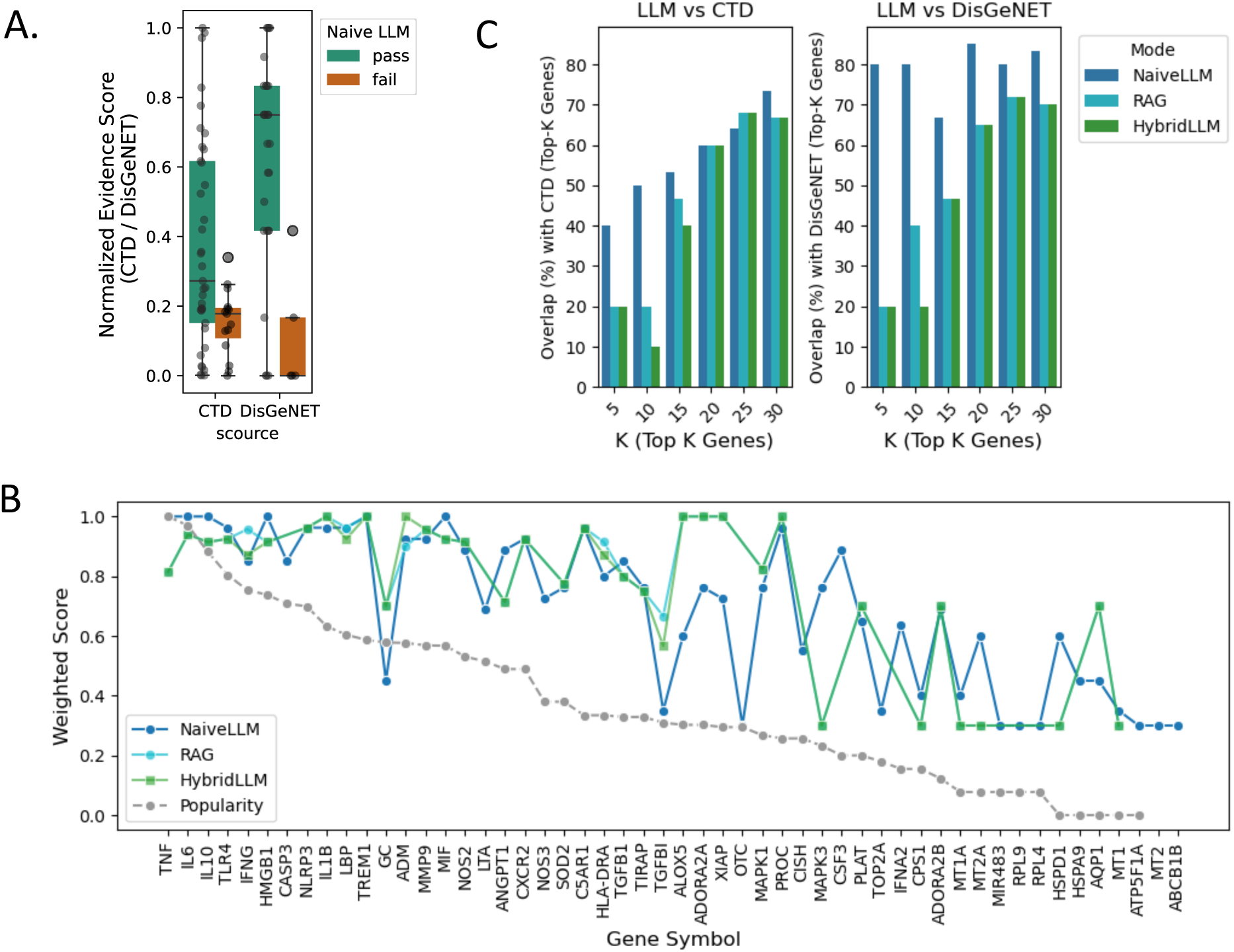
Validation of scoring methods on the 52-gene sepsis benchmark set **(A)** Normalized specificity scores for successfully identified versus missed genes across CTD and DisGeNET databases. Box plots show median, quartiles, and outliers, with individual data points overlaid. Successfully identified genes demonstrate significantly higher curation scores than missed genes, validating evidence-quality correlation. **(B)** Weighted scores across all 52 benchmark genes for each evaluation method compared to PubMed popularity (gray dotted line). Gene symbols are ordered by publication frequency. NaiveLLM (blue) shows strong correlation with evidence volume, while RAG (green) and HybridLLM (orange) exhibit more conservative scoring with occasional enhancements for understudied genes (e.g., AQP1, MIR483). All methods demonstrate appropriate evidence-weighted behavior. **(C)** Top-K overlap performance comparing method rankings against DisGeNET (left) and CTD (right) gene priorities. Y-axis shows percentage overlap between method top-K predictions and database top-K genes for K = 5, 10, 15, 20, 25, 30. NaiveLLM demonstrates superior ranking performance, particularly for top-5 predictions, while RAG and HybridLLM show lower immediate performance but retain capability for novel candidate detection. Performance patterns reflect method-specific trade-offs between ranking utility and literature verification.

Top-K overlap analysis revealed distinct method characteristics for practical applications (Figure 2C). NaiveLLM demonstrated superior ranking performance with 80% overlap for top-5 DisGeNET genes and 50% for CTD genes, maintaining strong performance through top-30 predictions. In contrast, RAG and HybridLLM showed substantially lower top-K performance, with only ∼20% overlap for top-5 predictions in both databases, improving gradually to 45-70% only at top-30. This pattern reveals a fundamental trade-off: while RAG and hybrid approaches sacrifice immediate ranking performance, they provide literature verification and enhanced detection of understudied candidates.

Systematic analysis identified 15 genes consistently missed across both databases, representing coherent biological categories including protein synthesis (RPL9, RPL4), metabolic regulation (OTC, CPS1, ATP5F1A, GC), cellular maintenance (MT1, MT2, MT1A, AQP1, ABCB1B, HSPA9, TGFBI), and regulatory mechanisms (MIR483, TOP2A). These categories primarily represent housekeeping functions that may be underrepresented in sepsis-specific literature despite biological relevance, highlighting the challenge of identifying functionally important but domain-understudied genes. This validation established framework reliability with characterized blind spots in metabolic pathways, providing performance boundaries essential for confident large-scale application.

### 3.2. Large-Scale Gene Screening and Functional Validation

Building on the validation results, we conducted genome-wide screening of the BloodGen3 repertoire (10,824 genes) to comprehensively identify sepsis-relevant gene candidates across the human transcriptome.

#### Genome-Wide Filtering and Prioritization

Genome-wide screening using our eight sepsis-related evaluation criteria (pathogenesis association, immune response, clinical biomarker potential, therapeutic relevance, and others) yielded 609 genes with >94% filtering efficiency (Figure 3A). We designated this filtered set as PS1 (Priority Set 1). To systematically analyze the score distribution, we stratified PS1 genes into five quantile-based groups using naive weighted scores: Q1 (top 100 genes), Q2 (109 genes), Q3 (144 genes), Q4 (120 genes), and Q5 (bottom 136 genes). This quantile-based stratification allows systematic examination of performance gradients across the priority gene set.

**Figure 3.**
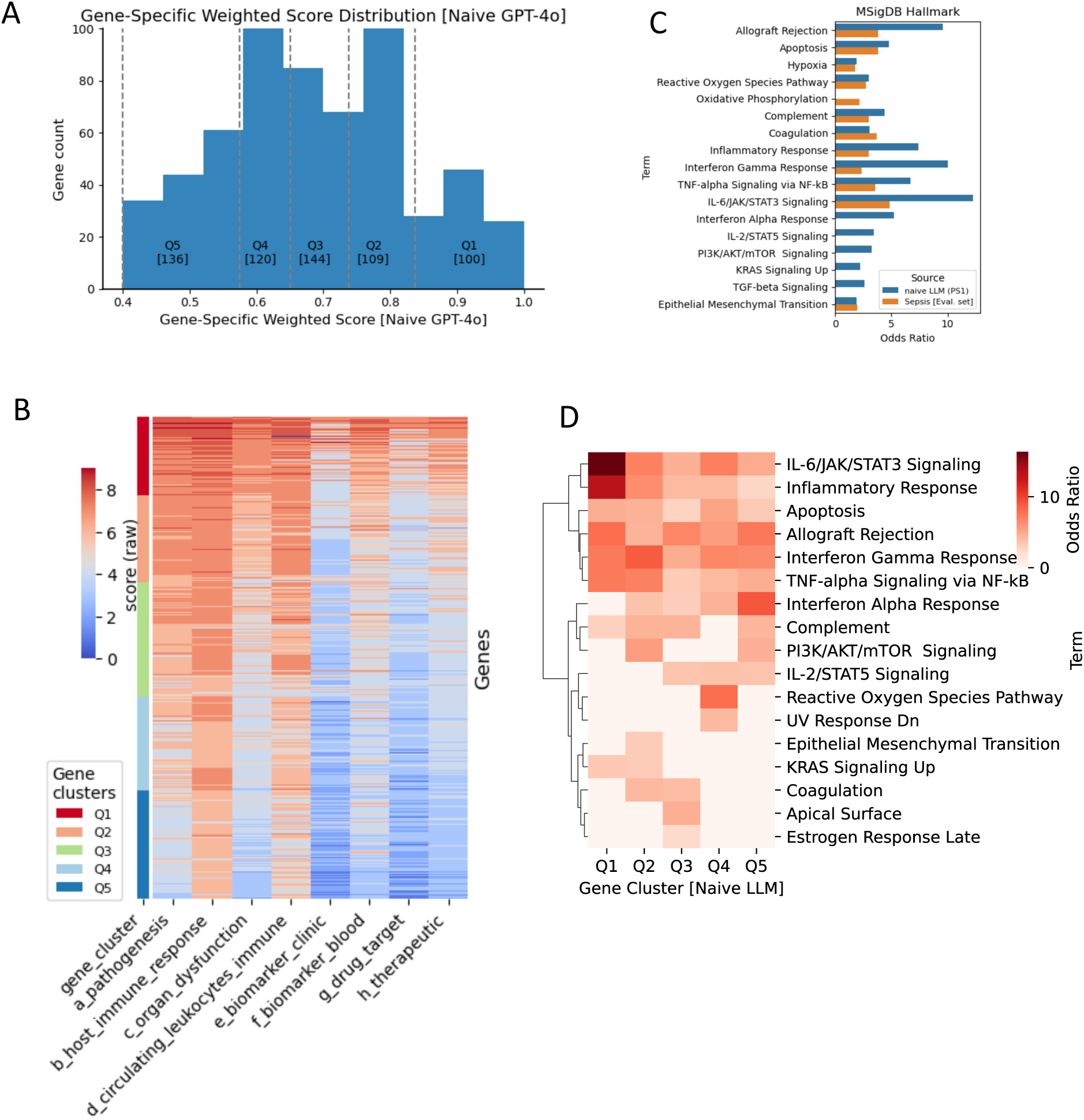
Naïve-LLM scoring of the BloodGen3 gene set and definition of Priority Set 1 (PS1) **(A)** Gene-specific weighted score distribution from naive LLM evaluation across 10,824 BloodGen3 genes. Histogram shows filtering efficiency with 609 genes (5.6%) meeting threshold criteria for PS1 (Priority Set 1). Score distribution demonstrates clear separation between selected and filtered genes. **(B)** Scoring heatmap for PS1 genes across eight sepsis-related evaluation criteria. Genes are stratified into quantile-based groups (Q1-Q5) ranked by naive weighted scores. Color intensity represents score magnitude, with Q1 genes (top quantile) showing consistently high scores across multiple criteria and lower quantiles exhibiting more heterogeneous patterns. **(C)** Functional enrichment analysis comparing PS1 genes against MSigDB Hallmark pathways. Bar plot shows odds ratios (log scale) for significantly enriched pathways (FDR < 0.05), with colors indicating gene set source. PS1 genes demonstrate strong enrichment for sepsis-relevant inflammatory and immune response pathways, validating biological coherence of the computational selection. Comparison with pooled evaluation gene set (gray bars) confirms maintained enrichment specificity. **(D)** Quantile-specific pathway enrichment heatmap showing functional stratification within PS1. Color intensity represents enrichment significance, with higher-scoring quantiles (Q1-Q2) showing stronger enrichment for core inflammatory pathways and lower quantiles exhibiting more diverse enrichment patterns. This gradient supports score-based prioritization validity.

#### Cross-Model Validation

To assess framework robustness across different LLM versions, we conducted comparative analysis between GPT-4o (current model) and GPT-4T (April 2024) on a subset of genes. The cross-model evaluation demonstrated high concordance with 74% agreement in gene prioritization decisions (Figure S6), indicating that our framework generates consistent results across model iterations and supporting the reliability of our large-scale screening approach.

#### Score Distribution Patterns Across Quantiles

The scoring heatmap revealed clear performance gradients across quantile groups (Figure 3B), as expected. Q1 genes (highest-scoring quantile) demonstrated consistently elevated scores across all eight evaluation criteria, representing the most promising sepsis candidates with strong evidence across multiple domains. Q2 genes maintained moderately high scores with some criteria-specific variations. The middle and lower quantiles (Q3-Q5) exhibited progressively more heterogeneous scoring patterns, with Q5 genes showing the most variable performance across criteria, suggesting these represent candidates with more limited or specialized evidence profiles.

#### Biological Coherence and Functional Stratification Analysis

To demonstrate that PS1 represents a biologically meaningful gene set rather than random selection artifacts, we performed functional enrichment analysis against MSigDB Hallmark pathways (Figure 3C, Table S4). The PS1 genes showed significant enrichment for pathways directly relevant to sepsis pathobiology, including TNF-α signaling via NF-κB, interferon gamma response, complement activation, inflammatory response, and reactive oxygen species pathways (all FDR < 0.05). This enrichment pattern confirms that our computational framework successfully captured genes with coherent biological relevance to the queried sepsis context. As a validation control, we compared PS1 enrichment patterns against the pooled evaluation gene set (929 genes) used in our initial framework development. The PS1 genes demonstrated comparable or enhanced enrichment magnitudes for sepsis-relevant pathways, confirming that the genome-wide screening successfully identified contextually appropriate genes rather than introducing systematic bias.

#### Score-Based Functional Stratification

Quantile-wise enrichment analysis revealed functional organization that correlates with our scoring hierarchy (Figure 3D). Higher-scoring quantiles (Q1-Q2) demonstrated stronger enrichment for inflammatory response pathways including IL-6/JAK/STAT3 signaling, complement activation, and TNF-α responses, consistent with their elevated scores across multiple sepsis-related criteria. Mid-tier quantiles (Q3-Q4) showed moderate enrichment for interferon responses and cellular stress pathways, while lower-scoring genes (Q5) exhibited more heterogeneous enrichment patterns including epithelial mesenchymal transition and metabolic processes. This functional gradient supports the validity of our scoring-based prioritization approach, suggesting that higher computational scores correspond to stronger associations with core inflammatory processes relevant to sepsis, while lower-scored genes may represent more peripheral or context-specific mechanisms. Consistent with evidence-based prioritization, PS1 gene clusters showed expected correlation with publication popularity (Figure S7), where higher-scoring clusters (Q1-Q2) contained genes with greater research attention compared to lower-tier clusters (Q4-Q5), validating that our framework appropriately weights established evidence while maintaining capability to identify less-studied but functionally relevant candidates.

This genome-wide screening successfully identified a functionally coherent set of sepsis candidates while maintaining high specificity for disease-relevant pathways and demonstrating robust cross-model consistency, providing a comprehensive foundation for subsequent detailed analysis.

### 3.3. Literature-Grounded Evaluation Through Retrieval-Augmented Generation

#### RAG Evaluation Framework and Performance

We implemented a sepsis-specific RAG system incorporating open source literature sources (reviews and research articles) and >10,000 PubMed abstracts using SPECTER 2 and LlamaIndex (see Methods). Each of the 609 PS1 genes was evaluated across eight sepsis-related queries, generating 4,872 total evaluation instances. RAG responses underwent independent faithfulness evaluation using LlamaIndex’s Faithfulness evaluator with Phi4 as an independent LLM model, providing binary Pass/Fail assessments based on alignment between retrieved literature chunks and generated justifications.

Of 4,872 queried instances, 1,484 passed the faithfulness test (30.5% pass rate), covering 455 unique genes designated as PS2 (Figure 4A). The stringent pass rate reflects the conservative nature of literature-grounded evaluation, ensuring high-confidence literature support for retained genes. Analysis of our scoring rubric categories revealed that high-scoring instances demonstrated the strongest correlation with faithfulness pass rates, validating our scoring criteria and confirming that computational confidence metrics appropriately reflect literature evidence quality.

**Figure 4.**
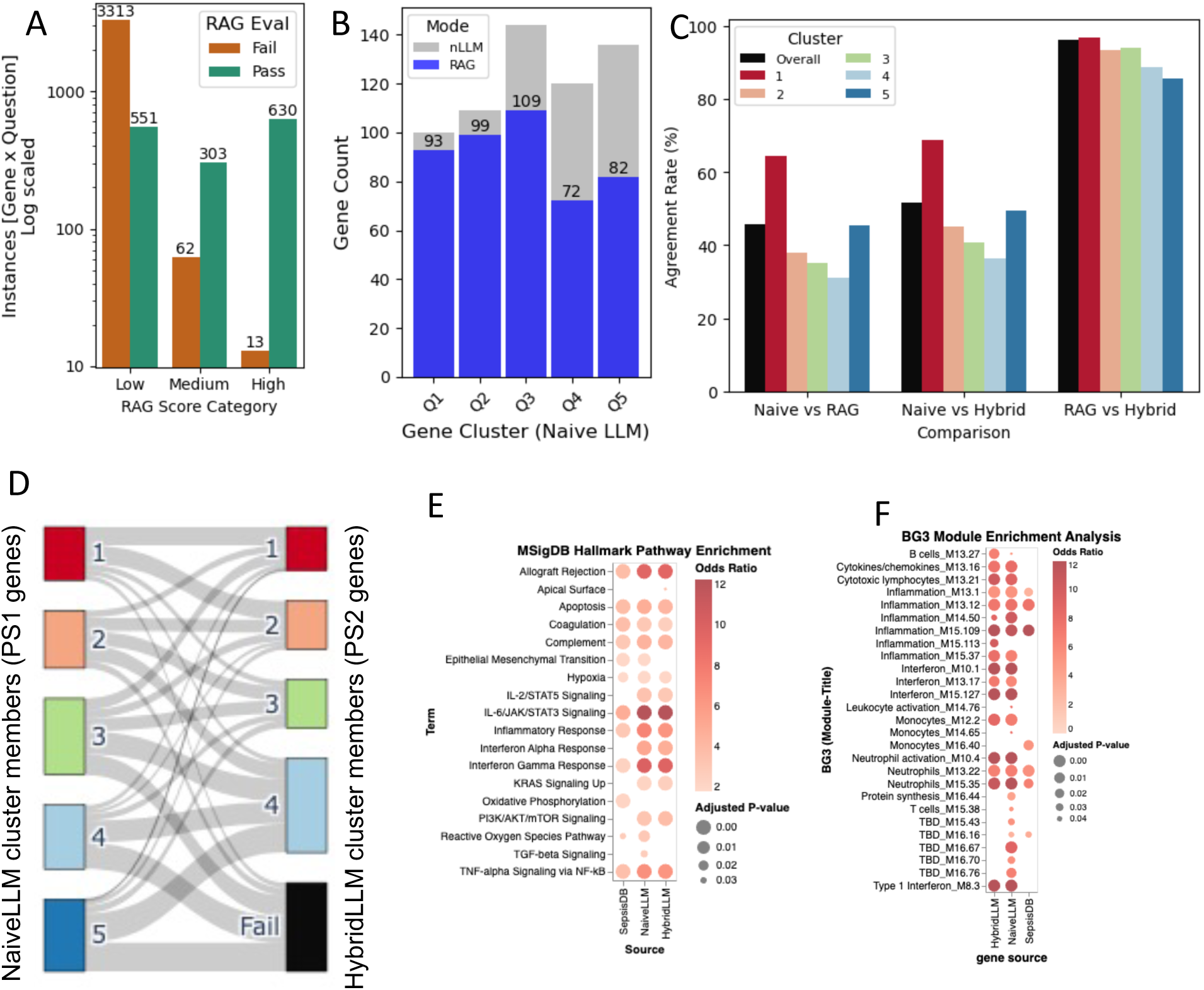
Retrieval-augmented evaluation and definition of Priority Set 2 (PS2) **(A)** RAG evaluation performance across 4,872 gene-query instances. Bar plot shows pass/fail distribution across RAG score categories (Low/Medium/High), with 1,484 instances passing faithfulness evaluation (30.5% overall pass rate). High-scoring instances demonstrate strongest correlation with literature validation success. **(B)** Gene recovery rates across PS1 quantile clusters following RAG evaluation. Bars show gene counts per cluster, demonstrating evidence-dependent recovery with highest rates in top-tier clusters (Q1: 93%, Q2: 90%) and progressive decline in lower-tier clusters. **(C)** Method agreement analysis comparing evaluation approaches. Left panel shows overall agreement percentages between method pairs. Right panels show cluster-wise agreement patterns, with extreme-scoring clusters (Q1, Q5) showing highest concordance and mid-tier clusters exhibiting more variable agreement reflecting literature context influence. **(D)** Sankey diagram visualizing cluster membership dynamics between NaiveLLM (PS1, left) and HybridLLM (PS2, right) approaches. Flow width represents gene count transitions, showing cluster consolidation from 5 to 4 clusters, with preserved high-confidence assignments and selective filtering of low-evidence genes. Colors correspond to cluster rankings. **(E)** MSigDB Hallmark pathway enrichment comparison across Evidence set (n=1,279), PS1 (n=609), and PS2 (n=442). Circle size represents adjusted p-values (<0.05), color intensity shows odds ratios. PS2 demonstrates enhanced enrichment for core sepsis pathways with selective loss of broader regulatory processes (TGF-β, EMT). **(F)** BG3 module enrichment analysis for PS2 genes showing cell-type specific signatures. Strong enrichment for inflammation, interferon response, monocyte activation, and neutrophil activation modules validates immune-focused biological coherence, with selective absence of non-specific gene modules confirming functional specificity.

#### Cluster-Wise Recovery and Evidence Stratification

RAG evaluation revealed expected evidence-dependent recovery patterns across PS1 clusters (Figure 4B). Top-tier clusters achieved highest literature validation rates: Q1 (93% recovery, 93/100 genes) and Q2 (90% recovery, 98/109 genes), consistent with their elevated baseline scores and extensive research documentation. Q3 showed moderate recovery (75%, 109/144 genes), while Q4 and Q5 both achieved approximately 60% recovery. This gradient confirms that higher computational scores correlate with stronger literature evidence availability, supporting the validity of our initial scoring framework (Figure S7).

#### Method Agreement and Scoring Dynamics

Comparative analysis between evaluation approaches revealed moderate but systematic agreement patterns (Figure 4C). Naive vs RAG comparison showed approximately 50% agreement, reflecting the conservative nature of literature-based filtering. Naive vs Hybrid demonstrated improved concordance, particularly for extreme-scoring genes where computational and literature evidence align. Cluster-wise agreement patterns showed expected conservation: Q1 exhibited maximum overlap due to consistently high-confidence literature support, while Q5 showed substantial agreement due to consistently limited evidence. Mid-tier clusters (Q2-Q4) displayed more variable agreement, indicating that literature context significantly influences evaluation outcomes for moderately evidenced genes (Table S6).

#### Re-clustering and Cluster Membership Dynamics

We re-clustered the 442 (filtered gene from RAG “pass” genes) using gene-wise weighted scores from HybridLLM, considering only the 1,484 RAG pass instances for scoring calculations. Following the same quantile-based clustering approach, we identified a maximum of 4 clusters for PS2, compared to 5 clusters in the original PS1 set, reflecting the more concentrated distribution of high-confidence, literature-validated genes.

The Sankey diagram reveals systematic shifts in cluster membership between NaiveLLM (PS1) and HybridLLM (PS2) approaches (Figure 4D). The flow patterns demonstrate several key dynamics: Cluster consolidation is evident as genes from PS1’s top clusters (Q1-Q2) predominantly flow into PS2’s highest-tier clusters, indicating preserved high-confidence assignments. Upward mobility is observed where some genes from PS1’s mid-tier clusters (Q2-Q3) migrate to higher PS2 clusters, suggesting that literature context enhanced their evidence profiles. Conservative filtering is apparent through the elimination of PS1’s lowest-confidence cluster (Q5), with these genes either being filtered out entirely or redistributed into PS2’s lower-tier clusters. The flow thickness patterns indicate that cluster retention is strongest for originally high-scoring genes, while cluster reassignment is most common for moderately scored genes where literature evidence significantly influenced final classification (Table S9).

#### Progressive Pathway Enrichment Through Filtering

Comparative functional enrichment analysis across the three gene sets reveals progressive enhancement of sepsis-relevant pathway signatures (Figure 4E). While the evidence set (n=927) shows broad pathway coverage with moderate enrichment intensities, both PS1 (NaïveLLM, n=609) and PS2 (HybridLLM, n=442) demonstrate substantially higher odds ratios for core inflammatory pathways including complement activation, TNF-α signaling, inflammatory response, and interferon pathways. This progressive enrichment intensity suggests successful noise reduction and signal enhancement through computational filtering and literature validation.

BG3 module enrichment analysis further validates PS2’s biological coherence by demonstrating specific enrichment for immune cell-type modules directly relevant to sepsis pathophysiology (Figure 4F). PS2 genes show pronounced enrichment for inflammation-associated modules, interferon response pathways, monocyte activation signatures, and neutrophil activation modules. This cell-type specificity aligns with established sepsis immunopathology, where monocyte/macrophage dysfunction and neutrophil activation are central disease mechanisms(31–34). Importantly, PS2 demonstrates selective absence of enrichment for non-specific gene modules, indicating successful filtering of housekeeping and broadly expressed genes that lack sepsis-specific relevance.

PS2 maintains enrichment for most core sepsis pathways present in PS1, while showing selective loss of broader regulatory pathways including TGF-β signaling and epithelial mesenchymal transition (EMT). This filtering pattern, also observed in SepsisDB-derived gene sets, reflects the literature-validation process’s preference for well-documented acute inflammatory processes over tissue remodeling and resolution mechanisms.

### 3.4. Multi-Dimensional Optimization Identifies High Confidence Priority Set 3 and Candidates genes

#### Cluster-Specific Functional Analysis and Final Prioritization

To identify the most promising therapeutic targets from PS2, we conducted cluster-specific functional enrichment analysis followed by multi-dimensional scoring optimization. Initial analysis of PS2’s four clusters using MSigDB revealed distinct functional specialization patterns (Figure 5A). The top performing cluster (Cluster 1) demonstrated the strongest enrichment across core sepsis pathways including allograft rejection, IL-6/JAK/STAT3 signaling, interferon gamma response, inflammatory response, and TNF-α signaling via NF-κB, establishing it as the highest-confidence functional cluster. We designated this gene set as Priority set 3 (PS3 n=82 genes: Table S9,S10).

**Figure 5.**
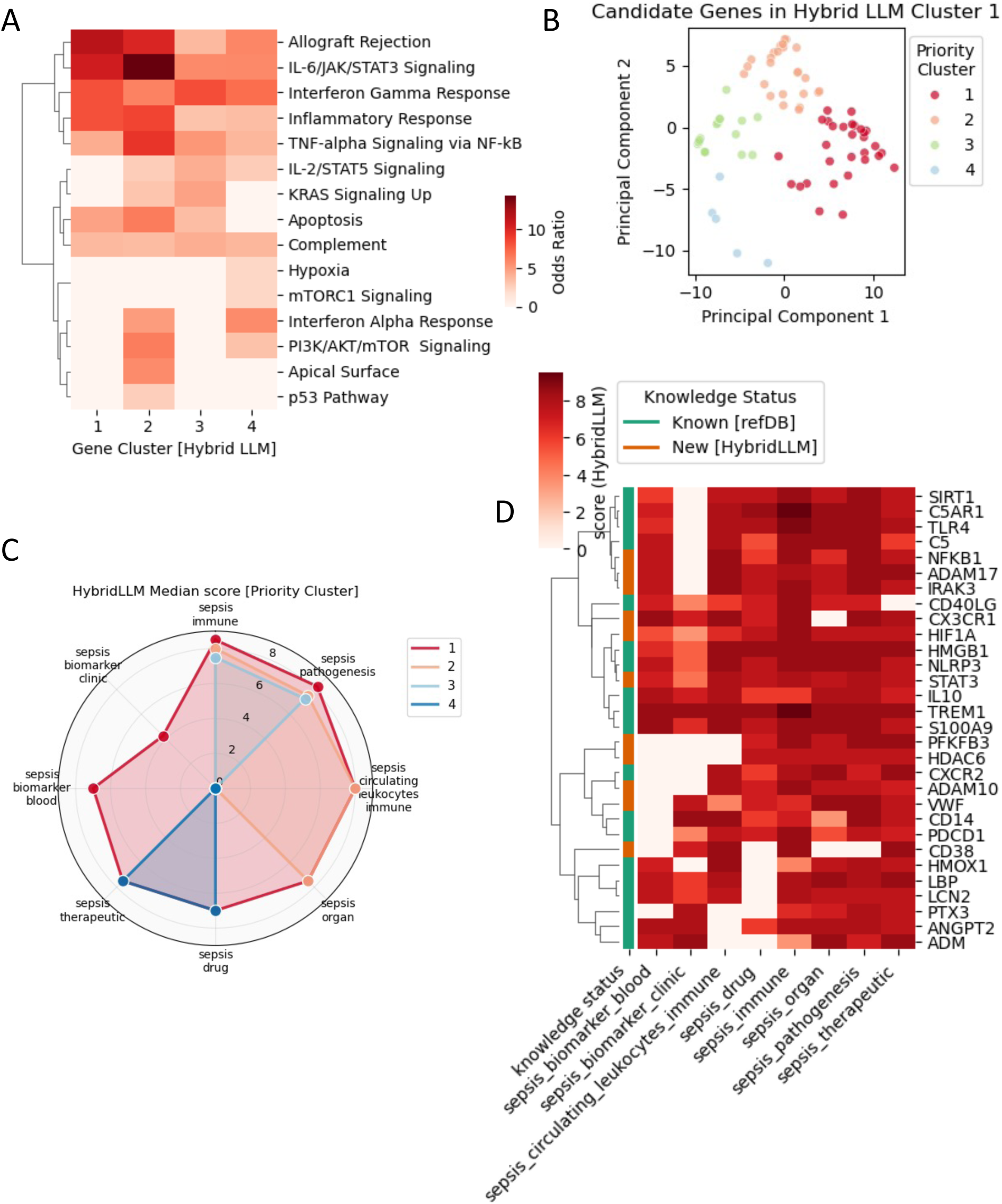
Principal-component optimization of PS2 and selection of Priority Set 3 (PS3) (A) Cluster-specific functional enrichment analysis for PS2’s four gene clusters using MSigDB Hallmark pathways. Heatmap shows odds ratios for significantly enriched pathways (FDR < 0.05), with Cluster 1 demonstrating strongest enrichment across core sepsis pathways including allograft rejection, IL-6/JAK/STAT3 signaling, inflammatory response, and TNF-α signaling via NF-κB. Color intensity represents enrichment magnitude. (B) Principal component analysis (PCA) of PS3 genes (n=82) based on HybridLLM scores across eight evaluation criteria. PCA reveals four distinct sub-clusters with diyerent scoring profiles. Points are colored by priority cluster assignment, with PC1 and PC2 explaining the major variance in multi-dimensional scoring space. (C) Polar plot comparison of mean HybridLLM scores across eight evaluation dimensions for the four PCA-derived clusters. Radar chart demonstrates that Priority Cluster 1 (red) consistently achieves highest scores across all criteria including pathogenesis association, immune response, biomarker potential, and therapeutic relevance, establishing it as the optimal candidate set. (D) Comprehensive scoring heatmap for candidate genes (n=30) selected from PCA Cluster 1. Shows HybridLLM scores across all eight evaluation queries with color intensity representing score magnitude (0-10 scale). Genes are annotated by discovery status: “Known” (from original evidence set) and “New” (novel candidates identified through computational pipeline). Both known and newly identified genes demonstrate consistently high scores across evaluation criteria, validating the framework’s ability to identify clinically relevant candidates while discovering novel targets with robust literature support.

#### Multi-Dimensional Score Analysis and Sub-Clustering

We selected PS3 for detailed multi-dimensional analysis using principal component analysis (PCA) across HybridLLM scores for all eight evaluation categories (Figure 5B). PCA revealed four distinct sub-clusters within this high-confidence gene set, indicating further functional or evidence-quality stratification. The PCA clustering captured genes with similar scoring profiles across the multi-dimensional evaluation space, allowing identification of the most consistently high-performing candidates. Comparative analysis using polar plots revealed distinct scoring profiles across the four PCA-derived clusters (Figure 5C). The radar chart comparison of mean Hybrid scores across eight evaluation dimensions demonstrated that PCA Cluster 1 genes consistently achieved the highest scores across all evaluation criteria, including pathogenesis association, immune response relevance, clinical biomarker potential, and therapeutic targeting feasibility. This comprehensive high performance across multiple independent evaluation dimensions established PCA Cluster 1 as the optimal candidate set (Table S10). This analysis established a clear hierarchy: PS2 (455 genes) → PS3 (82 genes from top cluster) → final candidate set (30 genes from optimal PCA cluster).

Based on the multi-dimensional optimization analysis of PS3 genes (n=82), we designated the 30 genes from the top-performing PCA as candidate set, representing high confidence sepsis genes. The comprehensive scoring heatmap reveals candidate set’s composition and validation status (Figure 5D, Table S10): genes are color-coded to distinguish between established candidates from our original evidence set (n=19 “Known”) and novel discoveries identified through our computational pipeline (n =11 “New”). All candidate genes demonstrate consistently high HybridLLM scores across the eight evaluation queries, with both known and newly identified genes showing comparable scoring profiles. Importantly, each candidate gene is backed by literature-grounded evidence from our RAG system and underwent additional verification through deep-search features to ensure robust evidence support (Supplementary Method 2).

#### Discovery and Validation Balance

The final candidate set successfully balances validation and discovery objectives by including both established sepsis genes that serve as positive controls and novel candidates that represent potential breakthrough targets. The comparable scoring patterns between known and new genes validates our computational framework’s ability to identify clinically relevant candidates while the literature grounding and deep-search verification ensures that novel discoveries possess substantial evidence support rather than representing computational artifacts.

This progressive filtering approach—from 10,824 genome-wide genes to 30 ultra-high confidence candidates—demonstrates successful integration of computational assessment, literature validation, and multi-dimensional optimization to identify the most promising sepsis therapeutic targets for immediate experimental investigation.

## 4. DISCUSSION

The benchmarking validation demonstrates systematic biological coherence through 71.2% recall with strong evidence quality correlation, distinguishing our approach from routine LLM applications that lack rigorous validation against expert-curated standards. RAG’s conservative behavior—frequent zero assignments with selective enhancement only when literature support exists—reflects appropriate uncertainty handling for biomedical inference where false discoveries incur significant costs. The systematic blind spots in metabolic pathways (protein synthesis, urea cycle, cellular maintenance) reveal coherent framework limitations that likely reflect literature emphasis on inflammatory over metabolic mechanisms in sepsis research, enabling informed large-scale result interpretation.

Recent biomedical LLM applications reveal fundamental limitations requiring sophisticated solutions. Kim et al. (2024) achieved only 16.0% accuracy with GPT-4 for phenotype-driven gene prioritization, significantly underperforming traditional bioinformatics tools(18). Recent RAG implementations demonstrate incremental improvements but lack comprehensive validation. GeneRAG (35) (2024) achieved 39% improvement in gene question answering through Maximal Marginal Relevance integration, while MIRAGE benchmarking showed medical RAG systems improving LLM accuracy by up to 18%(36). However, these approaches focus on general question-answering rather than systematic gene prioritization workflows. Wu et al. (2025) developed RAG-driven Chain-of-Thought methods for rare disease diagnosis from clinical notes, achieving over 40% top-10 accuracy, but without systematic faithfulness evaluation or biological validation(37). AlzheimerRAG (38) demonstrated multimodal integration for literature retrieval, and DRAGON-AI (39) applied RAG to ontology generation, but none address the core challenges of evidence verification and systematic validation in clinical contexts.

Early literature-based gene prioritization was established by Génie (2011), which ranked genes using MEDLINE abstracts and ortholog information but relied on basic text mining without modern NLP techniques (40). Established methods like ProGENI (16)(protein-protein interaction integration) and the Monarch Initiative(17) (curated database integration) demonstrate superior performance through structured knowledge but lack dynamic literature access and evidence verification capabilities that modern biomedical research demands.

Our framework addresses these limitations through systematic integration of curated domain knowledge (6,346 high-quality sepsis publications), dual-evaluator faithfulness assessment (71.94% inter-evaluator agreement), and Chain-of-Thought hybrid reasoning. The resulting 71.2% benchmark recall substantially exceeds recent LLM applications while maintaining biological coherence through systematic pathway enrichment validation.

The three-stage filtering approach successfully reduced computational noise while preserving biological signal, as evidenced by progressive pathway enrichment from PS1 (n=609) to PS2 (n=442) to PS3 (n=82). The 94% filtering efficiency demonstrates appropriate stringency, though the 30.5% RAG faithfulness pass rate may introduce bias toward well-studied genes. The final candidate gene composition (19 known, 11 novel candidates) with comparable scoring profiles supports balanced discovery-validation dynamics while avoiding computational artifacts.

Strong enrichment for sepsis-relevant pathways (TNF-α signaling, complement activation, interferon responses) confirms successful biological signal capture. The quantile-based functional stratification validates score-based prioritization, with higher-scoring clusters demonstrating stronger inflammatory pathway enrichment. However, selective loss of broader regulatory pathways (TGF-β signaling, epithelial mesenchymal transition) in literature-validated sets suggests potential under-representation of resolution mechanisms.

The 74% agreement between GPT-4o and GPT-4T versions provides moderate confidence in framework robustness across model iterations. This consistency level highlights the sensitivity of LLM-based approaches to model architecture differences. The systematic correlation between computational scores and PubMed publication frequency validates evidence-weighted behavior but reveals potential circularity favoring well-published genes.

The framework’s computational requirements present both advantages and limitations. Multi-stage evaluation requires substantial API costs and processing time that may limit accessibility. The dependency on proprietary models creates sustainability concerns for long-term research applications and reproducibility as models evolve.

Future iterations should incorporate experimental data beyond literature analysis, including functional genomics and clinical outcomes data. Integration of tissue-specific and temporal expression patterns could enhance context-relevant prioritization. Expanding beyond inflammatory mechanisms to include metabolic and repair pathways would provide more comprehensive target coverage.

The methodological framework provides a foundation for systematic AI-driven biomedical discovery with potential applications beyond sepsis research. However, continued validation against experimental outcomes and clinical data will be essential for establishing real-world utility and addressing the persistent challenge of translating computational predictions into therapeutic advances. The framework’s strength lies in systematic prioritization rather than definitive target identification, offering improved resource allocation for early-stage research while acknowledging the substantial experimental validation required for clinical translation.

## 5. Conclusions

We developed and validated a comprehensive computational framework for systematic gene prioritization in sepsis research, addressing critical limitations in current LLM-based biomedical applications. Through progressive filtering of 10,824 genes to 30 ultra-high confidence candidates, our approach demonstrates that rigorous methodology can transform unreliable LLM outputs into systematically validated biological insights. The framework’s key innovations include curated domain-specific knowledge base construction, dual-evaluator faithfulness assessment to reduce hallucination risks, and Chain-of-Thought hybrid reasoning that synthesizes computational predictions with literature evidence. Benchmark validation against expert-curated databases achieved 71.2% recall with strong evidence quality correlation, substantially exceeding recent LLM applications while maintaining biological coherence through systematic pathway enrichment. The identification of 11 novel high-confidence candidates alongside 19 established sepsis genes demonstrate balanced discovery-validation dynamics, though these candidates represent systematically prioritized hypotheses requiring extensive experimental validation rather than immediate therapeutic targets.

Our methodological approach—combining computational assessment, literature validation, and multi-dimensional optimization—provides a template for evaluating AI-driven biomedical discovery tools while establishing principles for responsible AI application in biomedical research. The modular design allows flexible adaptation where LLM models, gene lists, evaluation queries, and RAG databases can be systematically replaced to address any user-defined disease context or research question, extending applicability beyond sepsis to diverse biomedical domains. The systematic characterization of framework limitations, particularly blind spots in metabolic pathways and bias toward well-studied genes, enables informed interpretation of results and guides future improvements. While computational efficiency and model dependency present adoption challenges, this work demonstrates that sophisticated computational frameworks can bridge the gap between AI capabilities and clinically translatable applications, provided they incorporate rigorous validation, systematic bias characterization, and appropriate uncertainty handling. The framework’s strength lies in efficient resource allocation for early-stage research, offering a pathway for harnessing LLM potential in biomedical research while maintaining the scientific rigor essential for advancing from computational predictions to therapeutic breakthroughs.

## Reference

1. van ’t Veer LJ, Dai H, van de Vijver MJ, He YD, Hart AAM, Mao M, et al. Gene expression profiling predicts clinical outcome of breast cancer. Nature. 2002 Jan;415(6871):530–6.

2. Golub TR, Slonim DK, Tamayo P, Huard C, Gaasenbeek M, Mesirov JP, et al. Molecular Classification of Cancer: Class Discovery and Class Prediction by Gene Expression Monitoring. Science. 1999 Oct 15;286(5439):531–7.

3. Chaussabel D, Quinn C, Shen J, Patel P, Glaser C, Baldwin N, et al. A Modular Analysis Framework for Blood Genomics Studies: Application to Systemic Lupus Erythematosus. Immunity. 2008 Jul 18;29(1):150–64.

4. Bennett L, Palucka AK, Arce E, Cantrell V, Borvak J, Banchereau J, et al. Interferon and Granulopoiesis Signatures in Systemic Lupus Erythematosus Blood. J Exp Med. 2003 Mar 17;197(6):711–23.

5. Rinchai D, Altman MC, Konza O, Hässler S, Martina F, Toufiq M. Definition of erythroid cell-positive blood transcriptome phenotypes associated with severe respiratory syncytial virus infection. Clin Transl Med. 2020;

6. Rinchai D, Deola S, Zoppoli G, Kabeer BSA, Taleb S, Pavlovski I. High–temporal resolution profiling reveals distinct immune trajectories following the first and second doses of COVID-19 mRNA vaccines. Sci Adv. 2022 Nov;11;8(45):eabp9961.

7. Geiss GK, Bumgarner RE, Birditt B, Dahl T, Dowidar N, Dunaway DL, et al. Direct multiplexed measurement of gene expression with color-coded probe pairs. Nat Biotechnol. 2008 Mar;26(3):317–25.

8. Spurgeon SL, Jones RC, Ramakrishnan R. High throughput gene expression measurement with real time PCR in a microfluidic dynamic array. PloS One. 2008 Feb 27;

9. Rinchai D, Syed Ahamed Kabeer B, Toufiq M, Tatari-Calderone Z, Deola S, Brummaier T. A modular framework for the development of targeted Covid-19 blood transcript profiling panels. J Transl Med. 2020 Jul;31;18(1):291.

10. Altman MC, Rinchai D, Baldwin N, Toufiq M, Whalen E, Garand M. Development of a fixed module repertoire for the analysis and interpretation of blood transcriptome data. Nat Commun. 2021 Jul 19;

11. Li S, Rouphael N, Duraisingham S, Romero-Steiner S, Presnell S, Davis C, et al. Molecular signatures of antibody responses derived from a systems biology study of five human vaccines. Nat Immunol. 2014 Feb;15(2):195–204.

12. Chaussabel D, Pulendran B. A vision and a prescription for big data–enabled medicine. Nat Immunol. 2015 May;16(5):435–9.

13. Brummaier T, Kabeer BSA, Wilaisrisak P, Pimanpanarak M, Win AK, Pukrittayakamee S, et al. Cohort profile: molecular signature in pregnancy (MSP): longitudinal high-frequency sampling to characterise cross-omic trajectories in pregnancy in a resource-constrained setting. 2020 Oct 1 [cited 2025 May 30]; Available from: https://bmjopen.bmj.com/content/10/10/e041631

14. Rinchai D, Chaussabel D. A training curriculum for retrieving, structuring, and aggregating information derived from the biomedical literature and large-scale data repositories [Internet]. 2022. Available from: https://f1000research.com/articles/11-994

15. Abbas A, Rehman MS, Rehman SS. Comparing the Performance of Popular Large Language Models on the National Board of Medical Examiners Sample Questions. Cureus. 2024 Mar;16(3):e55991.

16. Emad A, Cairns J, Kalari KR, Wang L, Sinha S. Knowledge-guided gene prioritization reveals new insights into the mechanisms of chemoresistance. Genome Biol. 2017 Aug 11;18(1):153.

17. Putman TE, Schaper K, Matentzoglu N, Rubinetti VP, Alquaddoomi FS, Cox C, et al. The Monarch Initiative in 2024: an analytic platform integrating phenotypes, genes and diseases across species. Nucleic Acids Res. 2024 Jan 5;52(D1):D938–49.

18. Kim J, Wang K, Weng C, Liu C. Assessing the utility of large language models for phenotype-driven gene prioritization in the diagnosis of rare genetic disease. Am J Hum Genet. 2024 Oct 3;111(10):2190–202.

19. Singhal K, Azizi S, Tu T, Mahdavi SS, Wei J, Chung HW, et al. Large language models encode clinical knowledge. Nature. 2023 Aug;620(7972):172–80.

20. Deng L, Wang T, Yangzhang null, Zhai Z, Tao W, Li J, et al. Evaluation of large language models in breast cancer clinical scenarios: a comparative analysis based on ChatGPT-3.5, ChatGPT-4.0, and Claude2. Int J Surg Lond Engl. 2024 Apr 1;110(4):1941–50.

21. Chu Z, Chen J, Chen Q, Yu W, He T, Wang H, et al. Navigate through Enigmatic Labyrinth A Survey of Chain of Thought Reasoning: Advances, Frontiers and Future. In: Ku LW, Martins A, Srikumar V, editors. Proceedings of the 62nd Annual Meeting of the Association for Computational Linguistics (Volume 1: Long Papers) [Internet]. Bangkok, Thailand: Association for Computational Linguistics; 2024 [cited 2025 May 30]. p. 1173–203. Available from: https://aclanthology.org/2024.acl-long.65/

22. Toufiq M, Rinchai D, Bettacchioli E, Kabeer BSA, Khan T, Subba B. Harnessing large language models (LLMs) for candidate gene prioritization and selection. J Transl Med. 2023 Oct;16;21(1):728.

23. 23. Syed Ahamed Kabeer B, Subba B, Rinchai D, Toufiq M, Khan T, Yurieva M, et al. From gene modules to gene markers: an integrated AI-human approach selects CD38 to represent plasma cell-associated transcriptional signatures. Front Med. 2025;12:1510431.

24. Subba B, Toufiq M, Omi F, Yurieva M, Khan T, Rinchai D, et al. Human-augmented large language model-driven selection of glutathione peroxidase 4 as a candidate blood transcriptional biomarker for circulating erythroid cells. Sci Rep. 2024;14(1):23225.

25. Montani I, Honnibal M, Honnibal M, Boyd A, Landeghem SV, Peters H. explosion/spaCy: v3.7.2: Fixes for APIs and requirements [Internet]. Zenodo; 2023 [cited 2025 Jul 2]. Available from: https://zenodo.org/records/10009823

26. Piñero J, Ramírez-Anguita JM, Saüch-Pitarch J, Ronzano F, Centeno E, Sanz F, et al. The DisGeNET knowledge platform for disease genomics: 2019 update. Nucleic Acids Res. 2020 Jan 8;48(D1):D845–55.

27. Davis AP, Wiegers TC, Johnson RJ, Sciaky D, Wiegers J, Mattingly CJ. Comparative Toxicogenomics Database (CTD): update 2023. Nucleic Acids Res. 2023 Jan 6;51(D1):D1257– 62.

28. Gene Set Knowledge Discovery with Enrichr -Xie - 2021 - Current Protocols - Wiley Online Library [Internet]. [cited 2025 Jul 2]. Available from: https://currentprotocols.onlinelibrary.wiley.com/doi/10.1002/cpz1.90

29. Ochoa D, Hercules A, Carmona M, Suveges D, Gonzalez-Uriarte A, Malangone C, et al. Open Targets Platform: supporting systematic drug–target identification and prioritisation. Nucleic Acids Res. 2021 Jan 8;49(D1):D1302–10.

30. Krämer A, Green J, Pollard J Jr, Tugendreich S. Causal analysis approaches in Ingenuity Pathway Analysis. Bioinformatics. 2014 Feb 15;30(4):523–30.

31. van der Poll T, van de Veerdonk FL, Scicluna BP, Netea MG. The immunopathology of sepsis and potential therapeutic targets. Nat Rev Immunol. 2017 Jul;17(7):407–20.

32. Kwok AJ, Allcock A, Ferreira RC, Cano-Gamez E, Smee M, Burnham KL, et al. Neutrophils and emergency granulopoiesis drive immune suppression and an extreme response endotype during sepsis. Nat Immunol. 2023 May;24(5):767–79.

33. Bruserud Ø, Mosevoll KA, Bruserud Ø, Reikvam H, Wendelbo Ø. The Regulation of Neutrophil Migration in Patients with Sepsis: The Complexity of the Molecular Mechanisms and Their Modulation in Sepsis and the Heterogeneity of Sepsis Patients. Cells. 2023 Mar 24;12(7):1003.

34. Hotchkiss RS, Monneret G, Payen D. Sepsis-induced immunosuppression: from cellular dysfunctions to immunotherapy. Nat Rev Immunol. 2013 Dec;13(12):862–74.

35. Lin X, Deng G, Li Y, Ge J, Ho JWK, Liu Y. GeneRAG: Enhancing Large Language Models with Gene-Related Task by Retrieval-Augmented Generation [Internet]. bioRxiv; 2024 [cited 2025 May 30]. p. 2024.06.24.600176. Available from: https://www.biorxiv.org/content/10.1101/2024.06.24.600176v1

36. Xiong G, Jin Q, Lu Z, Zhang A. Benchmarking Retrieval-Augmented Generation for Medicine. In: Ku LW, Martins A, Srikumar V, editors. Findings of the Association for Computational Linguistics: ACL 2024 [Internet]. Bangkok, Thailand: Association for Computational Linguistics; 2024 [cited 2025 May 30]. p. 6233–51. Available from: https://aclanthology.org/2024.findings-acl.372/

37. Wu D, Wang Z, Nguyen Q, Wang K. Integrating Chain-of-Thought and Retrieval Augmented Generation Enhances Rare Disease Diagnosis from Clinical Notes [Internet]. arXiv; 2025 [cited 2025 May 30]. Available from: http://arxiv.org/abs/2503.12286

38. Lahiri AK, Hu QV. AlzheimerRAG: Multimodal Retrieval Augmented Generation for PubMed articles [Internet]. arXiv; 2024 [cited 2025 May 30]. Available from: http://arxiv.org/abs/2412.16701

39. Toro S, Anagnostopoulos AV, Bello SM, Blumberg K, Cameron R, Carmody L, et al. Dynamic Retrieval Augmented Generation of Ontologies using Artificial Intelligence (DRAGON-AI). J Biomed Semant. 2024 Oct 17;15(1):19.

40. Fontaine JF, Priller F, Barbosa-Silva A, Andrade-Navarro MA. Génie: literature-based gene prioritization at multi genomic scale. Nucleic Acids Res. 2011 Jul;39(Web Server issue):W455-461.

