## supplementary Information for "Automating Candidate Gene Prioritization with Large Language Models: From Naive Scoring to Literature-Grounded Validation"

### 1 Building Retrieval Augmented Generation (RAG) as a component of prioritization

#### 1.1 Knowledge Space Construction

A high-quality, curated knowledge base is essential for ensuring factual accuracy and relevance in a Retrieval-Augmented Generation (RAG) system. To build this knowledge space, we focused on open-access biomedical literature explicitly related to sepsis, applying both domain-specific and citation-based filters to maximize reliability.

##### 1.1.1 Document Retrieval and Curation

We queried NCBI PubMed Open Access using the advanced search interface to retrieve documents with "sepsis" in the title or abstract. To refine quality and ensure the documents were citation-relevant, we cross-referenced entries using OpenAlex (1) . The filtering criteria included:

- Open access availability
- High citation percentile, selecting only those with citation_n_pctile_value > 0.8
- This resulted in a final set of 6,346 documents, composed of: 4,441 original research articles and 1,905 review articles spanning from 1990 to 2025.

To enhance coverage, we supplemented this collection with 9,557 abstracts from relevant sepsis publications that were behind paywalls, ensuring comprehensive literature representation across both freely accessible and subscription-based sources.

##### 1.1.2 Literature Overview and Access Strategy

**Figure S1** provides a statistical overview of the curated literature:

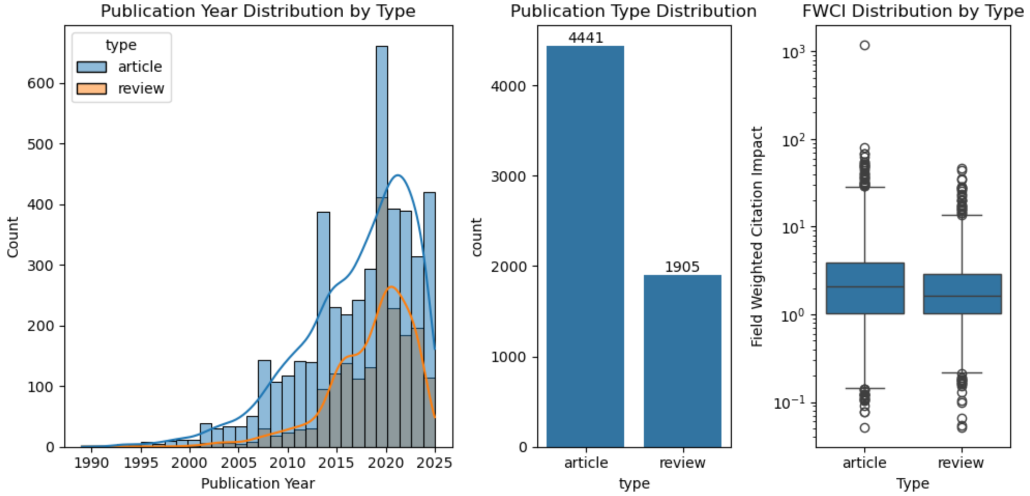

*Figure S1. Document corpus characteristics. Left: Publication year trends by article type; Center: Distribution of document types; Right: Field-weighted citation impact (FWCI), reflecting the quality of retained literature.*

We note that access limitations remain a challenge for full-text integration of paywalled research articles. To address this, we intentionally enriched the corpus with open-access review articles, which often synthesize findings from both open and proprietary sources. This strategy helped ensure thematic and temporal coverage without violating access restrictions.

#### 1.2 Preprocessing and Normalization

Each article was preprocessed to prepare it for downstream vectorization:

- Cleaning included removal of references, URLs, and non-informative characters.
- Segmentation was performed to chunk documents into paragraphs and sentence windows.
- Biomedical Named Entity Recognition (NER) was used to normalize gene names and biological concepts.

Metadata annotations were added, including publication year, type, and source. The result was a richly annotated, semantically coherent knowledge space optimized for embedding and retrieval.

#### 1.3 Vector Database Creation and Evaluation

Building on the curated knowledge space, we constructed a high-performance vector database to support semantic search and evidence retrieval in the RAG system. We use LlamaIndex for developing (https://github.com/run-llama/llama_index).

##### 1.3.1 Embedding Model: SPECTER2 for Biomedical Semantics

We used allenai/specter2_base (2), a transformer model trained on scientific citation graphs, to generate document embeddings. This model was chosen for its ability to capture scientific relevance and topic similarity, making it ideal for complex biomedical domains like sepsis. Each document chunk from the knowledge space was embedded into a high-dimensional vector representation suitable for similarity-based retrieval.

##### 1.3.2 Chunking and Tagging

Documents were parsed into overlapping chunks, enabling flexible retrieval resolution. Using SpaCy's biomedical NER (3), we extracted and tagged biological entities such as genes and gene products. Chunks without meaningful biomedical content were excluded to maintain retrieval precision. Each node was augmented with (a) Standard metadata: title, publication year, type, source and (b) Domain-specific tags: e.g., GENE_likeTags, enhancing filter and reranking capabilities

##### 1.3.3 Vector Indexing with ChromaDB

All processed nodes were stored in a persistent vector index using ChromaDB (https://github.com/chroma-core/chroma) . Embeddings were generated using SPECTER2 and inserted into ChromaDB in parallel. Each vector was indexed alongside its metadata and tag annotations for efficient retrieval and post-query filtering. This system enables both approximate nearest neighbor (ANN) search filtered retrieval based on metadata (e.g., publication type or year), as well as tag-based reranking, improving relevance and factual accuracy

#### 1.4 Limitations and Future Improvements

While the current vector database enables high-fidelity semantic retrieval, it is stored in a local persistent directory, requiring users to download the full index to access the embedded knowledge. This poses limitations in scalability, portability, and collaborative use—particularly in cloud-native or lightweight deployment scenarios. To overcome these constraints, future versions will support cloud-hosted vector databases, allowing for remote access, real-time updates, and seamless integration into distributed applications.

Moreover, our current use of rule-based named entity recognition (NER) captures surface-level biomedical terms but lacks domain-aware hierarchical resolution. In future releases, we plan to incorporate ontology-driven NER enhancements—leveraging curated biomedical knowledge bases such as UMLS (4), DisGeNET(5), or BioPortal(6). These systems will improve entity disambiguation, hierarchical classification, and concept linking, enhancing both retrieval accuracy and downstream interpretability. Together, these improvements will enable a more intelligent and extensible RAG system tailored for large-scale biomedical discovery.

### 2 Context Retrieval, Response Synthesis, and Faithfulness Evaluation

The effectiveness of Retrieval-Augmented Generation (RAG) depends not only on the quality of the indexed knowledge base, but also on the ability to retrieve relevant, compact, and non-redundant context in response to user queries. In our framework, we implemented a disciplined retrieval and synthesis strategy that prioritizes semantic precision, query isolation, and response verifiability.

To avoid context leakage and ensure interpretability, each gene-field instance was processed independently. For every gene and associated sepsis-related query, a mutually exclusive RAG cycle was executed, like our initial run with naïve llms. This design prevented cross-query contamination and ensured that the evidence presented to the language model (LLM) was specifically and solely relevant to the current task. As a result, each query-response pair remained self-contained and traceable.

Retrieval was initiated through a top-k vector similarity search using the vector database constructed from SPECTER2 embeddings. For each query, the vector index returned a ranked list of semantically proximal document chunks. These initial candidates were then subjected to a reranking stage using a cross-encoder model (SentenceTransformerRerank), which rescored the context passages based on their alignment with the query in a joint embedding space. This two-step process—coarse retrieval followed by fine reranking—ensured that the selected context was not only topically relevant but also interpretively matched to the question.

The highest-ranked context chunks were concatenated to form a prompt-specific evidence block. This block, along with the original query, was formatted into a structured prompt and passed to GPT-4o. The LLM was tasked with generating a score (on a 1–10 scale), a short natural-language justification for the score, and references corresponding to the original sources of the retrieved evidence. This approach provided both a decision and the rationale behind it, grounded explicitly in the retrieved literature.

To safeguard the factual integrity of these model-generated justifications, we implemented an automated faithfulness evaluation step. Using a lightweight scoring routine built on an auxiliary LLM, we evaluated whether the justification was supported by the retrieved evidence. Specifically, the FaithfulnessEvaluator (from LlamaIndex) compared the query, the retrieved context, and the generated response, and returned a binary label: “Pass” if the justification was faithful to the

evidence, or “Fail” if it introduced unsupported claims. Additionally, the evaluator provided textual feedback explaining the reasoning behind each judgment.
This post-hoc validation layer was critical in ensuring that model outputs remained faithful to the retrieved material. Combined with the rigorous retrieval strategy, query isolation, and interpretive reranking, this evaluation completed a robust pipeline for reliable and explainable gene relevance assessment in sepsis literature.

#### 2.1 Comparative Faithfulness Evaluation and Selection of Final Evaluator

To evaluate the reliability and interpretability of our RAG-generated justifications, we conducted a comparative faithfulness analysis using two independent evaluators: Phi-4 and GPT-o3-mini. This evaluation was performed on a substantial and targeted subset of our dataset—2,928 gene-query instances, derived from 175 genes in Cluster 1 and 191 genes in Cluster 5, each tested across eight standardized queries. Cluster 1 represents our biologically prioritized gene set, consistently scoring high in the naive model, while Cluster 5 was selected as a negative control, containing genes deprioritized by GPT-4T.

To ensure fair and representative sampling, we verified that each query topic was evenly represented across both clusters. As shown in Figure S2, the representative subset covered approximately 30–35% of total queries in each topic-cluster combination, confirming balanced coverage across thematic domains.

Figure S2. Percentage of gene-query instances included in the representative evaluation subset, stratified by cluster and question type.

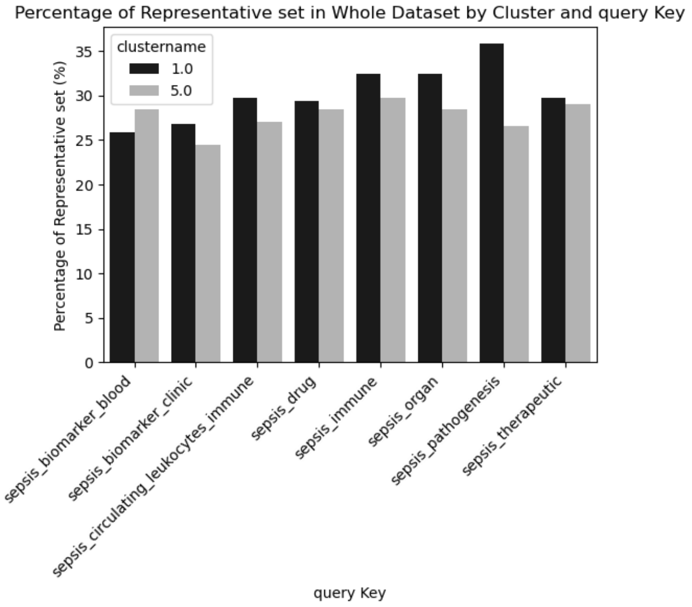

Each justification generated by GPT-4o was independently evaluated by Phi-4 and GPT-o3-mini, both of which returned a binary classification—“Pass” for responses grounded in the retrieved context, or “Fail” for those that were unsupported or hallucinated. The results of this dual-model evaluation are presented in Figure S3 and Table S1.

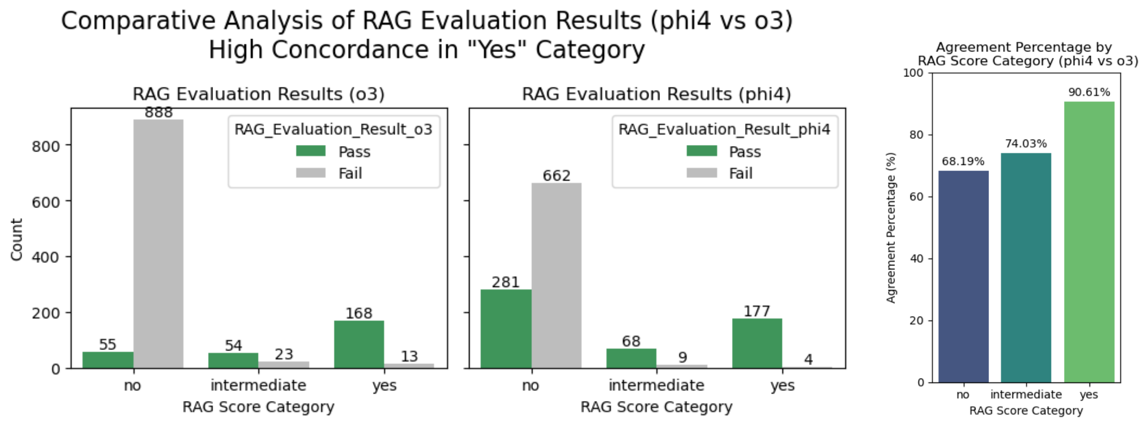

Figure S3. Comparative faithfulness evaluation using GPT-o3-mini and Phi-4. Left and center: Evaluation outcomes across RAG score categories. Right: Agreement percentages between the two evaluators per category.

We observed the highest agreement in the “yes” category, where 90.6% of RAG responses were rated consistently by both evaluators, while the overall agreement remains at 71.94% (Table S1). This finding highlights the reliability of the RAG system when strong contextual support exists. Disagreement was more frequent in the “no” category, where Phi-4 was more permissive (281 “Pass”) than GPT-o3-mini (55 “Pass”), likely due to differing thresholds for accepting negative assertions in the absence of explicit disconfirming evidence. The “intermediate” category showed moderate agreement, reflecting the interpretive ambiguity of these cases.

Given this comparative performance, we designated Phi-4 as our primary evaluator for the full dataset. This decision was based on the following justifications:

1. Evaluator independence: Phi-4 is architecturally and training-wise distinct from our generation model (GPT-4o), reducing evaluation bias.
2. Interpretive robustness: It consistently aligned with GPT-o3-mini in strong-evidence cases and provided stable judgments across score categories.
3. Scalability and reproducibility: Phi-4 offered lower variance in ambiguous cases, which was essential for applying this framework across all 2,925 gene-query instances.
4. Cost-efficiency and accessibility: Phi-4 was deployed locally using Ollama (https://github.com/ollama/ollama) , allowing high-throughput evaluation without incurring API or cloud-based inference costs. This also enhances reproducibility and enables use in resource-constrained settings.

Together, these criteria establish Phi-4 as a trustworthy, scalable, and reproducible evaluator for faithfulness assessment in large-scale RAG systems. Grounding our trust metrics in this model allows us to confidently propagate verified gene relevance predictions into further downstream biological analysis.

### 3 Prompts definitions:

#### 3.1 Queries

```
'sepsis_pathogenesis': "The gene {} is associated with the pathogenesis of sepsis.
Score: Based on evidence of the gene's involvement in the biological processes
underlying sepsis, including but not limited to its role in the dysregulated host
response to infection, organ dysfunction, or sepsis-related complications",

'sepsis_immune': " The gene {} is associated with the host immune response in sepsis.
Score: Based on evidence of the gene's involvement in the immune response during
sepsis, including but not limited to its role in innate or adaptive immunity,
inflammation, or immunosuppression",

'sepsis_organ': " The gene {} is associated with sepsis-related organ dysfunction.
Score: Based on evidence of the gene's involvement in the development or progression
of organ dysfunction in sepsis, including but not limited to its role in
cardiovascular, respiratory, renal, hepatic, or neurological dysfunction",

'sepsis_circulating leukocytes immune': " The gene {} is associated with the immune
response of circulating leukocytes in sepsis. Score: Based on evidence of the gene's
involvement in the immune response of circulating leukocytes during sepsis, including
but not limited to its role in leukocyte activation, migration, or function",

'sepsis_biomarker_clinic': " The gene {} or its products are currently being used as
a biomarker for sepsis in clinical settings. Score: Based on evidence of the gene or
its products' application as biomarkers for diagnosis, prognosis, or monitoring of
sepsis in clinical settings, with a focus on their validated use and acceptance in
medical practice",

'sepsis_biomarker_blood': " The gene {} has potential value as a blood
transcriptional biomarker for sepsis. Score: Based on evidence supporting the gene's
expression patterns in blood cells as reflective of sepsis or its severity,
considering both current research findings and potential for future clinical utility",

'sepsis_drug': " The gene {} is a known drug target for sepsis treatment. Score:
Based on evidence of the gene or its encoded protein serving as a target for
therapeutic intervention in sepsis, including approved drugs targeting this gene,
compounds in clinical trials, or promising preclinical studies",

'sepsis_therapeutic': " The gene {} is therapeutically relevant for managing sepsis
or its complications. Score: Based on evidence linking the gene to the management or
treatment of sepsis or its associated complications, including its role as a
potential target for adjunctive therapies or personalized treatment strategies"
```

#### 3.2 Prompt for naïve LLM response

```
You are an expert assistant in genomics and systems biology. Your task is to evaluate
the association between a specified gene and various biological or clinical aspects of
sepsis.

Please follow these instructions strictly:

1. Your answers must be precise, scientific, and based on high-quality evidence.
2. Use reputable sources including:
   - PubMed: https://pubmed.ncbi.nlm.nih.gov/
   - MGI: https://www.informatics.jax.org/
   - Google Scholar: https://scholar.google.com/
3. For each query, assign a score from 0 to 10 that reflects the strength of scientific
evidence supporting the gene’s involvement:
   - 0 – No evidence found
   - 1–3 – Very limited evidence
   - 4–6 – Some evidence, but needs validation or is context-dependent
   - 7–8 – Good evidence with consistent findings
   - 9–10 – Strong, well-supported evidence from multiple sources

4. Provide:
   - A score (integer 0–10)
   - A brief scientific justification
   - A list of scientific references that support your conclusion

5. Format your answer strictly as a JSON object using the following structure:
{
  "gene_name": "<gene_name>",
  "json_key": "<query_key>",
  "score": <integer 0–10>,
  "justification": "<concise scientific reasoning>",
  "references": ["<source 1>", "<source 2>", "..."]
}

Do not include any other explanation or text outside of the JSON object.
```

#### 3.3 Retrieval prompts

```
System:
    "You are a highly reliable scientific assistant.\n"
    "You must always reason step-by-step, following explicit roles.\n"
    "You must only output a valid JSON object as the final answer.\n"
    "Do not output anything outside the JSON structure."

User:
"You are a scientific research assistant. Your task is to evaluate the scientific
claim presented in the question using only the information provided in the
context.\n\n"
"Assign a score between 0 and 10, based on the strength of evidence in the
context:\n"
"- 0 = No supporting evidence\n"
"- 1–4 = Weak or indirect evidence\n"
"- 5–7 = Moderate or suggestive evidence\n"
"- 8–10 = Strong and direct evidence\n\n"
"Use only the information from the context. Do not use prior knowledge. If no
evidence is found, give a score of 0.\n"
"Include citations by listing the filenames of the documents that contain supporting
evidence.\n\n"
"Respond **only** with a valid JSON object. Do **not** include any explanations or
text outside the JSON.\n\n"
"The JSON response format is:\n"
"{\n"
"  \"score\": <integer from 0 to 10>,\n"
"  \"justification\": \"<brief explanation strictly based on the context>\",\n"
"}\n\n"
"Context:\n{context_str}\n\n"
"Question:\n{query_str}\n\n"
"Answer:"
```

#### 3.4 Hybrid prompt

```
You are an expert scientific assistant.

You must analyze a scientific question using both naive LLM knowledge and retrieved
scientific document evidence.

Question:
{query}

=========================================
Role 1: Naive LLM Critic
Think step-by-step:
- Analyze the Naive LLM evidence.
- How strong is the naive prior knowledge?
- Are there any weaknesses, biases, assumptions?
- How reliable would you consider it?

Evidence:
- Score (0-10): {naive_score}
- Justification: {naive_justification}

Write your reasoning before proceeding.

=========================================
Role 2: Retrieved Evidence Analyst
Think step-by-step:
- Analyze the Retrieved Context evidence.
- How specific, strong, or weak is the retrieved evidence?
- Are there conflicting findings?
- How much confidence can we place in it?

Evidence:
- Score (0-10): {rag_score}
- Justification: {rag_justification}

Write your reasoning before proceeding.

=========================================
Role 3: Final Arbiter
Think step-by-step:
- Compare the Naive LLM reasoning and Retrieved Context reasoning.
- If the Retrieved Context is strong and relevant, give it slightly more weight.
- If both sources agree, increase confidence.
- If they conflict, explain the discrepancy carefully.

At the end, output ONLY the following JSON format:

{{
  "final_answer": "<Yes/No/Unclear>",
  "final_score": <float between 0.0-10.0>,
  "scientific_explanation": "<your detailed scientific reasoning here, citing sources if
possible>"
}}
```

### 4 Benchmarking Strategy and Dataset Selection

#### 4.1 Rationale for Systematic Benchmarking

Automated gene prioritization methods in sepsis require systematic validation against curated biological knowledge to ensure reliability and clinical relevance (1,2). Unlike general text processing tasks, biomedical applications often lack definitive reference standards, particularly in sepsis—a multifaceted syndrome with heterogeneous molecular drivers (3). The complexity of sepsis pathophysiology, involving dysregulated host responses across multiple organ systems, necessitates evaluation against multiple independent knowledge sources to capture the full spectrum of gene-disease relationships (4). Furthermore, the clinical stakes of gene prioritization—where false negatives may overlook critical therapeutic targets and false positives may misdirect expensive validation efforts—demand systematic validation approaches that establish both sensitivity and specificity of the proposed methodology.

Effective benchmarking for sepsis gene prioritization must address several methodological considerations: the diversity of sepsis manifestations across patient populations, the evolution of sepsis definitions and diagnostic criteria over time, the integration of mechanistic knowledge with clinical observations, and the distinction between causal genes versus biomarker genes versus therapeutic targets. These considerations necessitate careful selection of reference datasets that provide clear curation methodologies, transparent evidence grading, and alignment with the specific aspects of sepsis biology under investigation.

#### 4.2 Available Sepsis Gene Datasets and Curation Methodologies

##### 4.2.1 DisGeNET: Expert-Curated Gene-Disease Associations

DisGeNET(5) represents one of the most comprehensive platforms for human gene-disease associations, integrating curated databases, text-mining results, and expert knowledge. For sepsis-related genes, DisGeNET employs a systematic curation approach combining manual literature review with natural language processing of biomedical texts. Each gene-disease association is assigned evidence scores based on the number and quality of supporting publications, with higher scores indicating stronger evidence from multiple independent sources. The sepsis gene set in DisGeNET (n=33) includes genes with direct experimental evidence for involvement in sepsis pathogenesis, immune response dysregulation, or clinical outcomes.

The curation methodology prioritizes peer-reviewed literature with experimental validation, clinical studies, and mechanistic investigations. DisGeNET's scoring system ranges from 0.0 to 1.0, with scores above 0.6 typically indicating strong literature support from multiple independent studies. The database provides transparent provenance tracking, allowing verification of evidence sources and assessment of confidence levels. For sepsis, the curated genes predominantly represent well-established players in innate immunity, inflammation, and organ dysfunction, making this dataset particularly suitable for validating the mechanistic aspects of sepsis gene prioritization (sepsis_pathogenesis, sepsis_immune, sepsis_organ queries).

##### 4.2.2 CTD: Chemical-Gene-Disease Interaction Networks

The Comparative Toxicogenomics Database (CTD) (6,7) provides manually curated associations between chemicals, genes, and diseases, with particular strength in therapeutic and toxicological relationships. CTD's sepsis gene set (n=52) is derived from systematic literature curation focusing on experimental evidence for gene involvement in sepsis through chemical interactions, therapeutic interventions, or toxicological mechanisms. Each association is supported by direct evidence from the literature, with curators manually extracting and verifying gene-disease relationships from peer-reviewed publications.

CTD's curation methodology emphasizes experimental validation and mechanistic understanding, with particular attention to genes that serve as targets for chemical interventions or demonstrate altered expression in response to sepsis-related stimuli. The database employs a numerical inference score for each gene-disease association, reflecting the strength of supporting evidence from curated literature. Additionally, CTD provides qualitative evidence descriptions and direct literature citations. CTD's strength lies in its focus on actionable gene-chemical-disease relationships, making it particularly valuable for validating therapeutic and drug target aspects of sepsis gene prioritization (sepsis_drug, sepsis_therapeutic queries). The systematic curation approach and emphasis on experimental evidence provide high-confidence associations suitable for benchmarking purposes.

##### 4.2.3 OpenTargets: Integrative Evidence Aggregation

OpenTargets (8) integrates multiple data sources including genetic associations, somatic mutations, drugs, pathways, and text mining to provide comprehensive gene-target-disease evidence. For sepsis, OpenTargets aggregates evidence from genome-wide association studies, expression quantitative trait loci, drug databases, and literature sources. The platform employs a sophisticated scoring system that weights different evidence types and combines them into overall association scores ranging from 0 to 1.

The platform focuses on gene targets that are at different stages of clinical trial development, which may result in a non-comprehensive representation of sepsis biology. The scoring methodology, while sophisticated, combines disparate evidence types that may not be equally relevant for specific aspects of sepsis biology. This heterogeneity makes OpenTargets valuable for comprehensive analysis but potentially problematic for focused benchmarking where clear ground truth is essential.

##### 4.2.4 MONARCH Initiative: Cross-Species Phenotype Integration

The MONARCH Initiative integrates phenotype and disease data across species, leveraging model organism research to inform human disease understanding (9). MONARCH's approach to sepsis involves mapping phenotypic annotations from model organisms to

human disease phenotypes through standardized ontologies. The platform includes genes associated with sepsis-like phenotypes in model organisms, immune dysfunction phenotypes, and inflammatory response abnormalities.
While MONARCH provides valuable cross-species insights, several factors limit its utility for focused sepsis benchmarking. The reliance on phenotype-to-disease mapping introduces uncertainty, as sepsis-like phenotypes in model organisms may not fully recapitulate human sepsis complexity. The database includes genes with varying degrees of evidence support, from well-characterized immunological players to genes with indirect phenotypic associations. Additionally, the cross-species integration may dilute human-specific sepsis mechanisms or include false positives from phenotypic similarities that do not reflect genuine mechanistic relationships.

##### 4.2.5 SEPON: SEPsis Ontology from NCBO BioPortal

SEPON (Sepsis Ontology) is a disease-specific ontology that defines the concepts and relationships between terms in the domain of sepsis and related diseases. It aims to support data standardization and enable precise, personalized diagnosis, treatment, and prognosis recommendations for sepsis. SEPON is hosted on NCBO BioPortal (access via: https://bioportal.bioontology.org/ontologies/SEPON), and was accessed programmatically via API for this analysis. However, no accompanying peer-reviewed literature describing its methodology or validation has been identified, limiting its immediate utility for rigorous benchmarking.

##### 4.2.6 IPA (Ingenuity Pathway Analysis): Commercial Knowledge Base

IPA (10) provides manually curated gene-disease associations derived from systematic literature review by professional scientific curators. The sepsis gene set in IPA (n=492) is substantially larger than other databases, reflecting broader inclusion criteria and commercial curation scope. IPA's curation process involves systematic review of peer-reviewed literature by trained scientists, with emphasis on mechanistic relationships and pathway-level associations.
However, several factors limit IPA's utility for transparent benchmarking. The commercial nature of the database restricts access to detailed curation methodologies, evidence scoring systems, and provenance information. The large gene set size may reflect less stringent inclusion criteria compared to more focused academic databases. Additionally, the lack of transparent scoring or confidence metrics makes it difficult to stratify associations by evidence quality, potentially including genes with weak or indirect evidence. In contrast, DisGeNET and CTD offer full transparency on curation and evidence scoring, reinforcing their suitability as benchmarking datasets.

#### 4.3 Gene Set Overlap Visualization

To further support the rationale behind benchmarking dataset selection, we evaluated the overlap among sepsis-related gene sets from six major resources: DisGeNET, CTD, OpenTargets, MONARCH, SEPON, and IPA. To empirically illustrate how these resources align, we visualized the overlap of sepsis gene sets from six sources (Figure 1). The heatmap underscores the concentrated agreement between DisGeNET and CTD, compared to the broader and more diffuse content in platforms like IPA and SEPON.

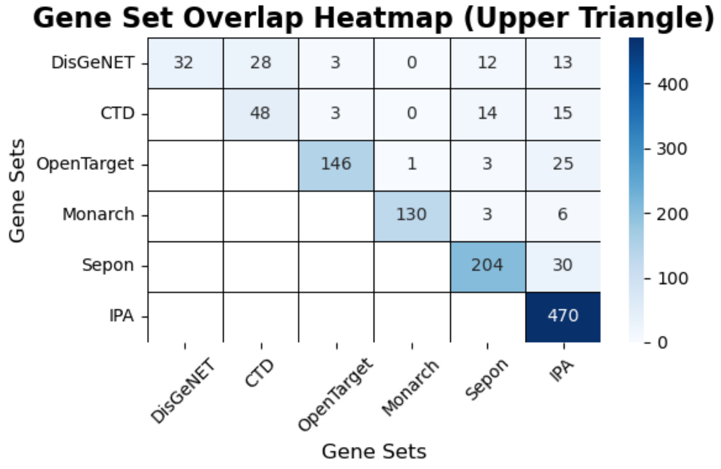

**Figure S4. Gene Set Overlap Heatmap**: This heatmap illustrates the intersection of gene sets across six resources related to sepsis research (n=929). The diagonal cells represent the total unique genes in each resource, while the off-diagonal cells show the shared genes between pairs of resources. The visualization highlights the extent of agreement and complementarity among the datasets. Darker shades represent higher gene overlap.

#### 4.4 Benchmarking Dataset Selection Rationale

Consideration of Additional Resources: Several exploratory resources were reviewed during dataset selection, including BIDOS (http://www.swmubidos.com/), SC2sepsis (11), SeptiSearch(12), MetaSepsisBase(13), and GeneCards (14). These platforms offer valuable complementary insights, such as gene expression dynamics, single-cell specificity, and curated biomarker annotations. However, they were excluded from benchmarking due to limitations in evidence traceability, lack of systematic curation, or absence of peer-reviewed documentation. For example, BIDOS and SC2sepsis aggregate omics data but lack standardized confidence scoring for gene-disease associations. SeptiSearch and MetaSepsisBase provide enrichment analysis and curated gene sets, but do not specify inclusion thresholds or validation standards. These resources remain useful for downstream validation or enrichment analysis, but do not meet the rigor required for defining gold-standard benchmarking sets.

Based on the methodological assessment of available datasets, we selected DisGeNET and CTD as primary benchmarking datasets for several critical reasons:

**Methodological Transparency:** Both databases provide clear documentation of curation processes, evidence requirements, and inclusion criteria, enabling transparent evaluation of benchmarking validity. The systematic literature-based curation approaches align with evidence-based medicine principles and provide traceable provenance for each gene-disease association.

**Evidence Quality Standards:** DisGeNET and CTD employ rigorous evidence standards requiring experimental validation, peer-review publication, and mechanistic understanding. This contrasts with databases that include computational predictions, automated text mining results, or indirect associations without experimental support.

**Complementary Coverage:** DisGeNET's strength in mechanistic and pathophysiological associations complements CTD's focus on therapeutic and chemical interaction relationships, together covering the primary aspects of our eight-query framework (mechanistic: sepsis_pathogenesis, sepsis_immune; therapeutic: sepsis_drug, sepsis_therapeutic).

**Appropriate Dataset Size:** The moderate sizes of DisGeNET (n=33) and CTD (n=52) sepsis gene sets enable detailed validation analysis while representing manageable, high-confidence reference standards. This contrasts with either very small datasets lacking statistical power or very large datasets potentially diluted with lower-confidence associations.

The combination of DisGeNET and CTD provides a robust, transparent, and methodologically sound foundation for evaluating our automated gene prioritization pipeline, establishing both the sensitivity to recover known sepsis genes and the biological coherence of the underlying methodology. Hence, these two data bases were compiled as "gold - standards", that we will use for our benchmarking. Other genes with causal relation and inferred association will be treated as reference for functional enrichment analysis (along with gold standards).

### 5 Distribution of LLM scores across question prompts

Supplementary Figure S5: Distribution of LLM-assigned scores across evaluation criteria before and after filtering

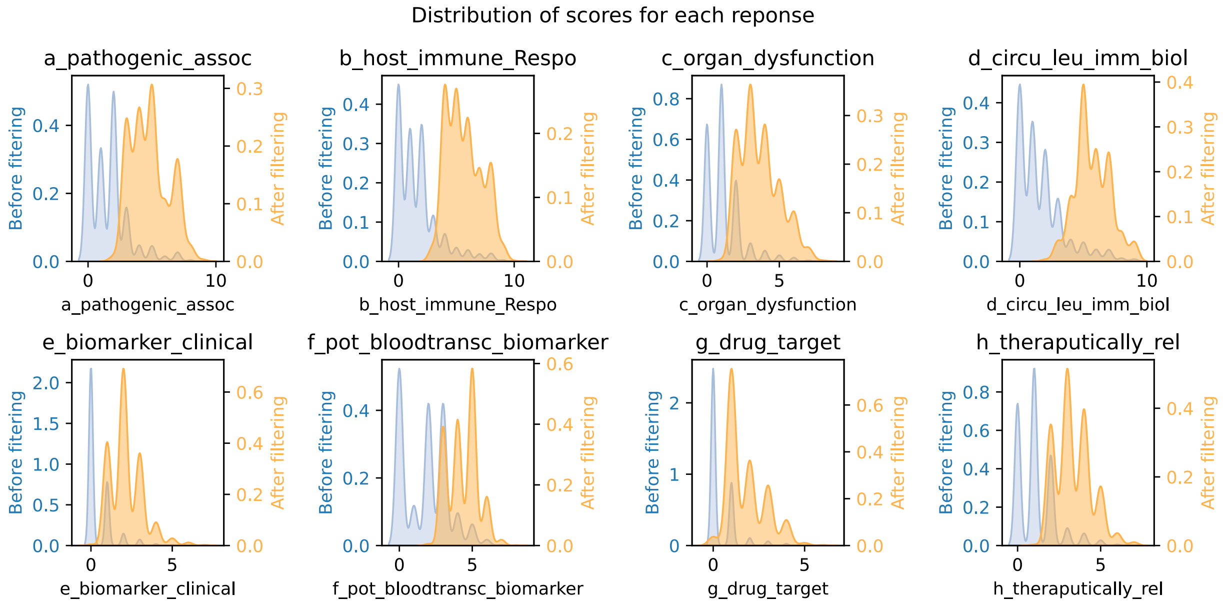

Figure S5: Distribution of scores assigned by GPT-4T for each of the eight evaluation criteria (a-h) in sepsis-related gene prioritization. Each panel represents one criterion: (a) association with pathogenesis of sepsis, (b) association with host immune response, (c) association with sepsis-related organ dysfunction, (d) relevance to circulating leukocytes immune biology, (e) current use as a clinical biomarker, (f) potential value as a blood transcriptional biomarker, (g) known drug target, and (h) therapeutic relevance. Blue distributions show scores across all genes in the BloodGen3 repertoire (n=10,824). Orange distributions show scores for the filtered subset of genes that scored > 5 in at least one criterion (n=1,070). The right-skewed patterns, particularly pronounced in criteria 'e' and 'g', reflect the biological reality where only a subset of genes have strong documented evidence for specific functions or clinical applications. Y-axis shows density of genes, and X-axis shows the score range (0-10). The density scales are shown on the left (blue, before filtering) and right (red, after filtering) of each panel.

### 6 Cross-Model Consistency Assessment

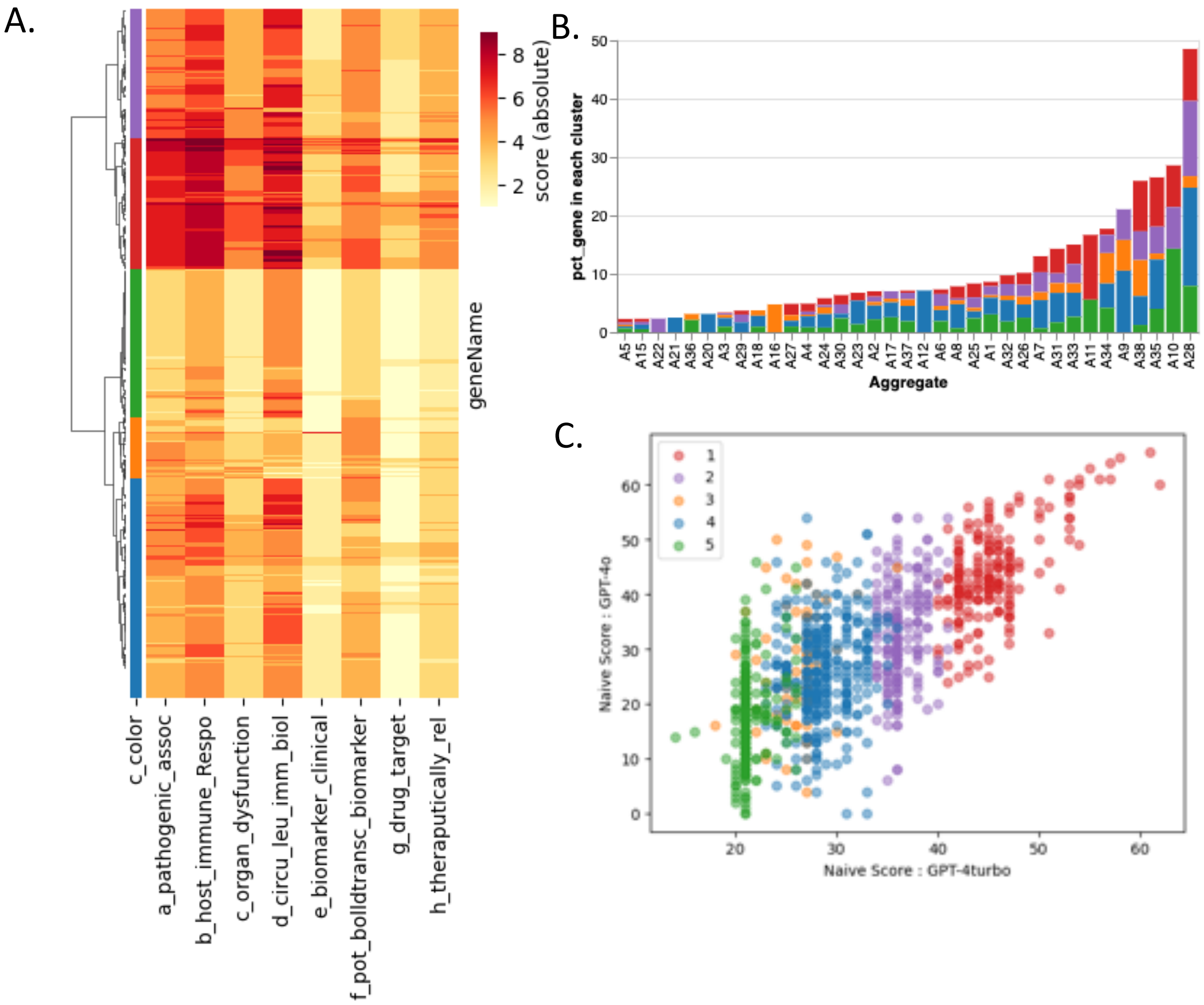

Figure S6. Naïve LLM performance in GPT-4T and 4o.

[A] Hierarchical Clustering of Sepsis-Related Genes Based on Response Scores : This figure presents a hierarchical clustering analysis of 1,070 genes (rows) filtered based on their relevance to sepsis-related queries (i.e genes with score > 5 in at least one of the sepsis related query). The heatmap displays the absolute response scores for each gene, with colors ranging from white (low scores) to orange to red (high scores). Genes are grouped into clusters based on their overall response patterns, as indicated by the color-coded track along the hierarchical clustering tree.

[B] Percentage of Gene in each cluster across Module Aggregates: Bar plot shows distribution of sepsis related genes (n=1070) across pre-defined BloodGen3 Module Aggregates (= 35 [out of 38]). Each bar shows percentage of genes found in each Module Aggregate, as well proportionality of gene clusters with same color code as shown in A.

[C] Correlation of Naive Relevance Scores Between GPT-4T and GPT-4o Across Clusters: Scatterplot comparing naive gene relevance scores assigned by GPT-4T (x-axis) and GPT-4o (y-axis) across all gene-query instances. Each point represents an individual instance, color-coded by cluster (1–5). The strong positive correlation across all clusters indicates broad semantic consistency between the two models. However, distinct vertical stratification—particularly visible in lower-ranked clusters (e.g., Cluster 5, green)—highlights differences in score distribution and scale compression in GPT-4T. In contrast, GPT-4o shows smoother gradation, suggesting improved contextual calibration. This visualization confirms that while both models share overall prioritization trends, GPT-4o offers more nuanced differentiation within and across clusters.

### 7 Gene popularity and filtered gene set by LLM for Sepsis

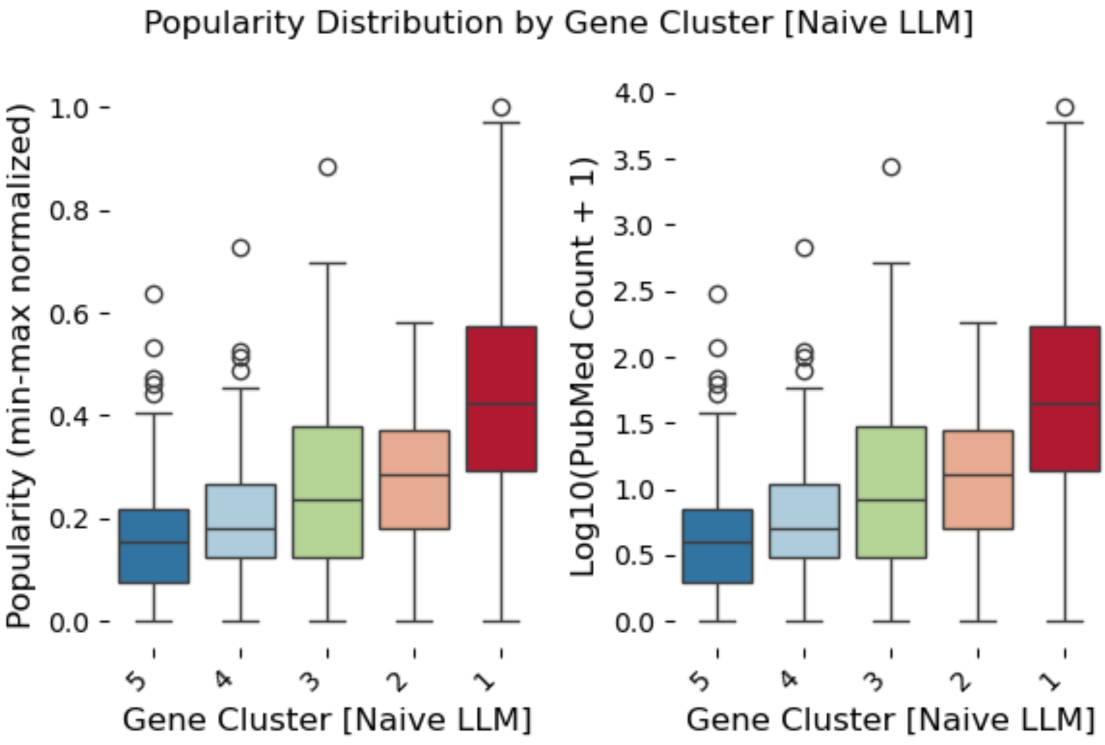

Figure S7 : Popularity distribution across gene clusters demonstrates expected correlation between computational scores and publication frequency, validating evidence-based prioritization approach..

### 8 Supplementary Tables

Table S1. Comparison of GPT-4T and GPT-4o Scores with Cluster and IPA/LLM Evidence Labels: This table lists all gene-query pairs evaluated in the study, including raw scores from GPT-4T and GPT-4o, their corresponding categorical interpretations (“no”,“yes,” “Intermediate,” ), the cluster assignment of each gene, and its evidence classification (LLM-only, IPA-only, or both). The table provides a comprehensive view of how gene relevance scores shift across LLM versions and how these relate to curated (IPA) versus generative (LLM) knowledge support.

Table S2: contingency Table of Naïve LLM runs GPT-4T vs GPT-4o.

Table S3: Classification report of naïve runs GPT-4T / GPT-4o
A: classification report of GPT-4T vs GPT-4o as per cluster assignment by initial run by GPT-4T.
B: Comparison w.e.t assigned evidence category (IPA)

Table S4: Comparing Faithfulness of RAG retrieval with Phi4 and GPT-o3-mini

Table S5: Contingency tables of Naive-RAG-Hybrid (A: Naïve Vs RAG; B: Hybrid Vs Naïve C: Hybrid Vs RAG)

Table S6: Classification Report of Naive-RAG-Hybrid Hybrid (A: Naïve Vs RAG; B: Hybrid Vs Naïve C: Hybrid Vs RAG)

Table S7: List of gene prioritized for high confidence scope (final score : Yes)

Table S8: List of gene prioritized for broad scope (final score Yes+ Intermediate)

Table S9: Detail result on 2928 (gene-query) instance with naïve scores, rag scores with retrieved nodes text, justification (by phi4), retrieval status , final score justification and hybrid scores.

### 9 Supplementary Figures

Figure S1: Distribution of LLM-assigned scores across evaluation criteria before and after filtering

Figure S2. Document corpus characteristics. Left: Publication year trends by article type; Center: Distribution of document types; Right: Field-weighted citation impact (FWCI), reflecting the quality of retained literature.

Figure S3. Percentage of gene-query instances included in the representative evaluation subset, stratified by cluster and question type.

Figure S4. Comparative faithfulness evaluation using GPT-o3-mini and Phi-4. Left and center: Evaluation outcomes across RAG score categories. Right: Agreement percentages between the two evaluators per category.

### 10 Reference

1. Priem, J., Piwowar, H., & Orr, R. (2022). OpenAlex: A fully-open index of scholarly works, authors, venues, institutions, and concepts. ArXiv. https://arxiv.org/abs/2205.01833 - Google Search [Internet]. [cited 2025 May 29].

2. Singh A, D'Arcy M, Cohan A, Downey D, Feldman S. SciRepEval: A Multi-Format Benchmark for Scientific Document Representations. In arXiv; 2022 [cited 2025 May 29]. Available from: https://arxiv.org/abs/2211.13308

3. Neumann M, King D, Beltagy I, Ammar W. ScispaCy: Fast and Robust Models for Biomedical Natural Language Processing. Proc 18th BioNLP Workshop Shar Task. 2019;319–27.

4. Bodenreider O. The Unified Medical Language System (UMLS): integrating biomedical terminology. Nucleic Acids Res. 2004 Jan 1;32(suppl_1):D267–70.

5. Piñero J, Ramírez-Anguita JM, Saüch-Pitarch J, Ronzano F, Centeno E, Sanz F, et al. The DisGeNET knowledge platform for disease genomics: 2019 update. Nucleic Acids Res. 2020 Jan 8;48(D1):D845–55.

6. Whetzel PL, Noy NF, Shah NH, Alexander PR, Nyulas C, Tudorache T, et al. BioPortal: enhanced functionality via new Web services from the National Center for Biomedical Ontology to access and use ontologies in software applications. Nucleic Acids Res. 2011 Jul 1;39(suppl_2):W541–5.
